# Convex-PL^*R*^ – Revisiting affinity predictions and virtual screening using physics-informed machine learning

**DOI:** 10.1101/2021.09.13.460049

**Authors:** Maria Kadukova, Vladimir Chupin, Sergei Grudinin

## Abstract

Virtual screening is an essential part of the modern drug design pipeline, which significantly accelerates the discovery of new drug candidates. Structure-based virtual screening involves ligand conformational sampling, which is often followed by re-scoring of docking poses. A great variety of scoring functions have been designed for this purpose. The advent of structural and affinity databases and the progress in machine-learning methods have recently boosted scoring function performance. Nonetheless, the most successful scoring functions are typically designed for specific tasks or systems. All-purpose scoring functions still perform poorly on the virtual screening tests, compared to precision with which they are able to predict co-crystal binding poses. Another limitation is the low interpretability of the heuristics being used.

We analyzed scoring functions’ performance in the CASF benchmarks and discovered that the vast majority of them have a strong bias towards predicting larger binding interfaces. This motivated us to develop a physical model with additional entropic terms with the aim of penalizing such a preference. We parameterized the new model using affinity and structural data, solving a classification problem followed by regression. The new model, called Convex-PL^*R*^, demonstrated high-quality results on multiple tests and a substantial improvement over its predecessor Convex-PL. Convex-PL^*R*^ can be used for molecular docking together with VinaCPL, our version of AutoDock Vina, with Convex-PL integrated as a scoring function. Convex-PL^*R*^, Convex-PL, and VinaCPL are available at https://team.inria.fr/nano-d/convex-pl/.

## Introduction

Rigorous estimation of the binding free energy requires exhaustive sampling in a certain thermodynamical ensemble. However, this approach is computationally prohibitive for virtual screening applications. Therefore, multiple attempts have been made to approximate the binding free energy by minimizing or totally excluding the sampling step. This resulted in a considerable number of scoring functions, ^1–31^ which, in general, aim to approximate the free energy change upon binding. The binding Gibbs free energy can be written as ^32,33^

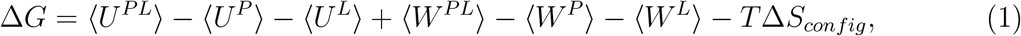

where the *P* superscript refers to the interactions with the protein, *L* - with the ligand, ⟨*U*⟩ and ⟨*W*⟩ are the averaged potential and solvation energies, respectively, and Δ*S*_*config*_ is the entropy change related to protein and ligand motions upon complex formation. However, many of the approaches would make very crude approximations of the entropic term and interactions with the solvent in the above equation. This causes the known flaw of many knowledge-based scoring functions preventing them from being used in screening tests. More precisely, many of them have a strong bias toward bigger and tighter protein-ligand interfaces. Conformations of a ligand inside a binding pocket that have a higher number of interactions with the protein, even weak ones, will often be preferred over the native ligand pose. However, in reality, some parts of the binding site and the ligand exposed to the solvent could be more favorable compared to the corresponding protein-ligand contacts.

The preference of larger interfaces can be illustrated by the publicly available results of scoring functions evaluation on the virtual screening test of CASF-2013^34^ and CASF-2016^35^ benchmarks shown in Figure 1 (a). Here one can see that the majority of the assessed scoring functions prefer binding with non-native ligands (decoys) with, on average, up to twice bigger buried solvent-accessible surface area (SASA) values than a native ligand has. Figure 1 (b) shows that, in fact, for some scoring functions this trend is even stronger if the total number of atoms is used instead of the SASA, buried upon binding. Notably, AutoDock Vina^9^ and AutoDock Vina-based Δ_Vina_RF_20_ ^22^ scoring functions do not suffer from this bias that much. This can be explained by the way AutoDock Vina scales its binding energies by the number of ligand’s rotatable bonds. GlideScore-XP^7^ also does not express any considerable bias toward the overall ligand molecule size neither. This is probably owing to its solvation term and the correct penalization of contacts between polar and hydrophobic groups.

**Figure 1:**
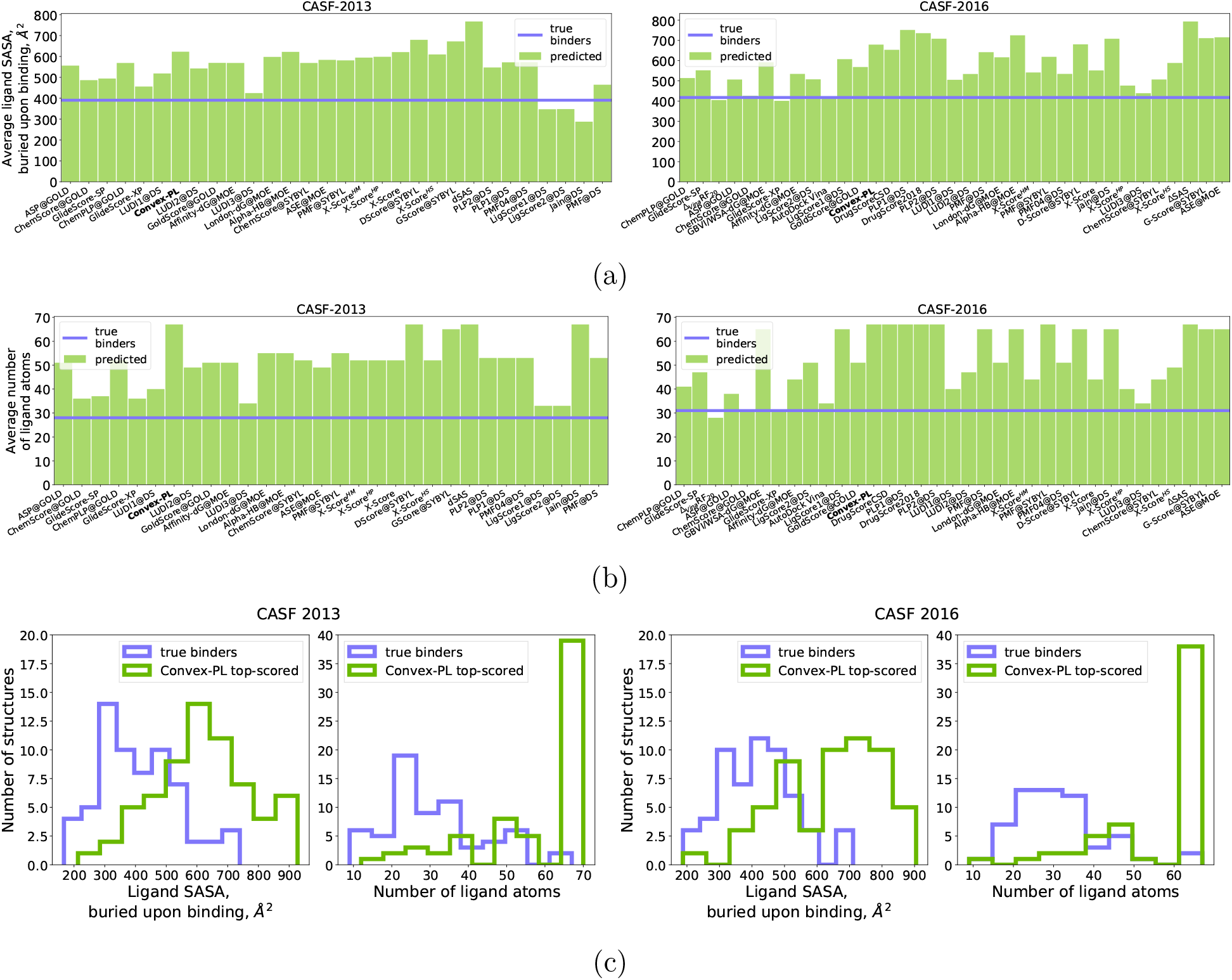
(a)-(b) The purple line represents the average values of buried SASAs and numbers of atoms computed for ligands that natively bind target proteins and should have been predicted as the most affine binders^[1]^. The green boxes correspond to decoys that were top-ranked by scoring functions assessed on the virtual screening test from the CASF benchmarks 2013 and 2016. The scoring functions are sorted by the ability to predict the highest affinity binder in the 5% of the top-ranked decoys. (c) Histograms of average ligand buried SASA and number of atoms computed for the native “truly binding” ligands (purple) and decoy poses, top-ranked by Convex-PL (green) in the virtual screening tests from the CASF-2013 and CASF-2016 benchmarks. SASA values were computed with PyMOL’s^36^ *get area()* function with *dot solvent* set to 3. ^[1]^ Or be among the most affine binders in several cases when the target protein was known to bind ligands with higher affinity but without co-crystal structure.

Many other empirical^5,6,9,26,37^ and some knowledge-based^8,15,38^ scoring functions circumvent these problems by including additional entropic and solvation terms in their expressions. A classical approximation of ligand conformational entropy is the number of torsions or atoms involved in rotatable bonds.^5,6,9,15,38,39^ More rigorous estimations may include sampling of the ligand conformational space. ^26,40^ Some scoring functions also include rigid-body contributions approximated with a logarithm of the ligand mass,^5,15^ even though this approximation, as well as the involvement of the mass-dependent rigid-body entropy itself, is arguable.^33,41,42^ Basic implicit representations of solvation include interaction terms proportional to the SASA,^22,38,43^ or solvent-accessible volume difference upon binding.^44–47^ Some algorithms utilize SASA in more sophisticated ways, such as calibration using the octanol-water partition coefficients alongside with separate hydrogen bonds description aiming at a better hydrophobic effects representation,^48,49^ or integrating the surface curvature factor of the molecules over the solvent-accessible surface area.^50^ Another way to compute solvation energy change with an implicit solvent model is to use the 3D-RISM,^51^ Poisson-Boltzmann, and generalized Born methods.^32,52–54^ They are, in general, much more computationally demanding, although some approximate solutions are applied in virtual screening. ^55^ Explicit solvent representation for molecular docking purposes^26^ requires either high-quality X-ray structures with the hydration shell resolved or the hydration shell sampling performed by dedicated algorithms and molecular dynamics-based pipelines.^56–59^ Although these approaches look quite intuitive and generally improve the docking quality, they are mainly used to predict the water sites for individual targets, which may require manual intervention, making it hard to apply them on a larger scale. However, this problem can be solved using statistical potentials for water molecules prediction.^60,61^

Despite the variety of possible estimations of entropic and solvation terms, Figure 1 implies that some of these strategies are yet not sufficient. In particular, our knowledge-based scoring function called Convex-PL^16^ demonstrated excellent results in the pose prediction tests but turned out to perform rather average in virtual screening exercises, as illustrated in Figure1 (c). Indeed, we can see that SASAs of about half of the ligands predicted to be the best binders are considerably higher than those of the true binders. This clearly indicates a bias of Convex-PL towards larger binding interfaces. The current work revisits the derivation of empirical scoring functions and presents a reworked Convex-PL model. Firstly, we retrained it on a more diverse structural dataset containing cofactors and modified residues to improve the support of interactions between ligands and molecules other than standard amino acids. Then, motivated by the general thermodynamics principles, we augmented Convex-PL with several physics-based terms that aim to penalize the bias of large interfaces. While the inclusion of the entropic terms in most cases had a clear enhancing effect, the incorporation of solvation terms turned out to be more challenging and, for Convex-PL, lead to controversial results. Throughout the manuscript, we will keep referring to the initial version of Convex-PL as to Convex-PL, the newer version, trained with the *regression* model, will be called Convex-PL^*R*^. Convex-PL^*R*^ does not require any specific preparation and parametrization of input structures. It is designed for relatively small molecules of less than 100 heavy atoms. Although we support ligand-cofactor interactions, the representation of metals is still limited. Currently, Fe has the most trustworthy representation driven by its presence in a considerable amount of heme cofactors. Although the dataset also included Zn, Mg, and other metals, we are not fully confident in them.

## Model

Following the protein-ligand binding free energy derivation in the canonical ensemble suggested in Gilson et al. ^33^, the *standard binding free energy* of a protein *A* and a ligand molecule *B* in a solvent can be written as

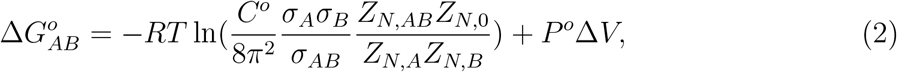

where *C*^*o*^ and *P*^*o*^ are the standard concentration and pressure, *σ*_*A*_, *σ*_*B*_, *σ*_*AB*_ are the symmetry numbers of each molecule, *Z*_*N,AB*_, *Z*_*N,A*_, *Z*_*N,B*_ are the configurational integrals of the protein-ligand complex, ligand, and protein in a solvent, respectively, *Z*_*N*,0_ is a configurational integral of this solvent containing *N* atoms, and Δ*V* is a solute volume change upon binding. The integration in the partition functions is taken over the internal coordinates **r**_*A*_, **r**_*B*_, **r**_*S*_ of the receptor, ligand and solvent, respectively, and six external coordinates *ζ*_*B*_ of the ligand molecule B that are defined relative to the protein molecule A.

Our general aim is to approximate the configurational integrals ratios *using only a single conformation* of the protein-ligand complex instead of rigorously sampling the conformational ensemble. Firstly, we suppose that the ratio 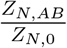 of the complex and the solvent configurational integrals can be approximated with the original Convex-PL knowledge-based potential,^16^ which can be represented as a dot product between a *structure* vector **x** and a *scoring* vector **w**,

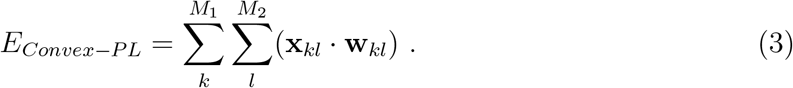

In this equation, the Convex-PL potential is considered to be a sum of pairwise interactions between the protein and the ligand atoms of *M*_1_ and *M*_2_ atom types, correspondingly. The *structure vector* **x** consists of number density functions extracted from the structures of protein-ligand complexes, and the *scoring* vector **w** contains weights for each contribution that are optimized by solving a *classification convex* optimization problem on a training dataset. The Convex-PL derivation is described in more detail in the corresponding paper.^16^ It is important to note, however, that the given form of the Convex-PL function implicitly contains some additional interactions, especially the hydrophobic ones associated with the solvent entropy change.

The ligand configurational integral ratio can be rewritten as follows:

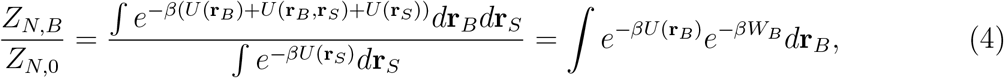

where *U*(**r**_*B*_), *U*(**r**_*S*_), *U*(**r**_*B*_, **r**_*S*_) are the potential energy of the ligand, solvent, and the interactions between the ligand and solvent, respectively, and the integration is taken over all internal coordinates of the ligand *B* and solvent *S*. Also, *W*(*B*) is the ligand *solvation energy* expressed as

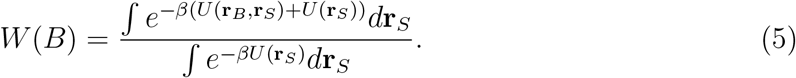

We assume that for *sufficiently small ligands*, whose initial local geometries were minimized by some conformer generator, we can neglect the intra-ligand interactions, *U*(**r**_*B*_) = 0, over the sampled ligand conformations *d***r**_*B*_. We also presume that in this case, *W*_*B*_ is *almost constant* in the sampled volume. Thus, we obtain the following expression,

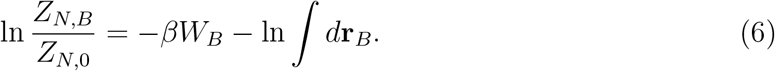

We approximate the first term with a set of descriptors containing solvent-accessible surface areas of atoms, and grid-based descriptors representing the displaced solvent volume that are described in more detail in the section below. The second term corresponds to the volume of the ligand conformational space, which we approximate with the logarithm of the number of ligand conformational states that can be adopted by rotations about *rotatable bonds*,

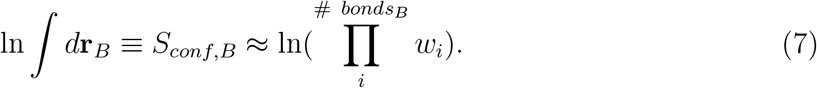

Here, the product is taken over the ligand rotatable bonds, and the weights *w*_*i*_ = 4 *− b*_*i*_ are computed using *i*th bond order *b*_*i*_. We only consider those bonds, for which the buried solvent-accessible surface area of at least one atom is greater than zero. The conformational symmetry is partially taken into account by not counting bonds with the terminal atoms.

A similar procedure can be carried out for the receptor configurational integral, resulting in

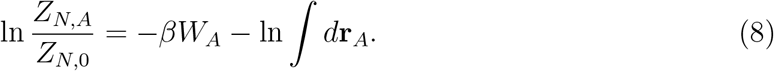

Following Gilson et al. ^33^, we can split the integration coordinates *d***r**_*A*_ to the *interface pocket* and *rigid* parts of the protein and neglect the integral over the rigid one. In general, the interface pocket integral can then be taken over the rotameric states of the pocket residues,^40,41,62,63^ however, we decided to approximate it with the volumes of hemispheres that a single residue can adopt by rotations around its C*α*–C*β* bond. The proportion of pocket residue’s bonds with allowed rotations in the unbound state can be estimated by the fraction of its solvent-accessible surface area *s*_*i,unbound*_ in the unbound state, and the total surface area of the same residue, if it is extracted from the receptor, *s*_*i,total*_. Thus, we obtain

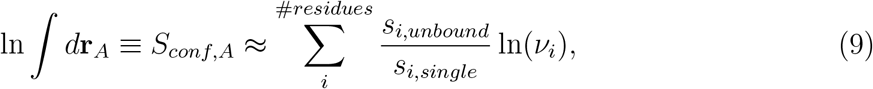

where the sum is taken over all the residues at the protein-ligand interface, and *v*_*i*_ are constant rotational volumes, precomputed for each amino acid. Neglecting the displaced volume, constants, and some of the symmetry effects, we get our final equation,

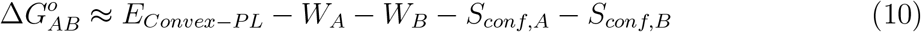

### Interactions with the solvent

As we have mentioned above, Convex-PL implicitly counts some interactions with the solvent in the corresponding pairwise potentials. To account for the potentially underestimated interactions in the *W*_*A*_, and *W*_*B*_ terms, we collected two types of descriptors. The first one was the buried solvent-accessible surface areas of atoms of different types that we computed using the POWERSASA library^64,65^ with a 1.4 Å probe atom radius. The second one is described in more detail in SI. It was based on statistics collected for the distance distributions between the grid points that represented solvent, and the atoms of the complex, protein, and ligand of different types.

After model training and ablation studies, it turned out that our solvent contributions do not have a significant effect. When we considerably regularize them, solvent contributions provide a subtle improvement on the CASF benchmarks, but almost no changes on other tests – and thus can be neglected. Alternatively, without a considerable regularization, weights of the solvent contributions trained to predict binding affinities started spoiling the virtual screening performance. This effect can be, in principle, circumvented by multi-task learning,^19,24,30^ but we decided to use a model without additional solvent terms, assuming that a portion of desolvation effects is implicitly included in the Convex-PL’s statistics.

### Regression model with entropic contributions

Convex-PL was initially trained to discriminate native and non-native ligand poses by solving an optimization problem, in which native and non-native poses within each protein-ligand complex are compared to each other to minimize the prediction loss. Comparing poses of *different* complexes to each other would be meaningless since the ‘native’ and ‘non-native’ class labels are relevant *inside* each complex only. This method is very efficient for reconstructing a scoring function for pose prediction. Its design, however, limits the ability to predict the absolute binding affinities. Nonetheless, we can circumvent this problem using regression towards available affinity data. Therefore, we combined the binding constants, the Convex-PL energy, and flexibility descriptors in a regression model, which lead to a considerable improvement of the CASF benchmark screening test results. We have tried solving both linear and non-linear kernel and ensemble regression problems with inclusion and exclusion of solvent features, and settled upon the linear ridge regression model. It is easily regularizable, fast to train, and interpretable. Table S5 of SI lists the resulting feature weights.

### Training data

We were using two separate datasets for the classification and regression parts of the model. Complexes used in the two CASF benchmarks were excluded from both datasets.

The original Convex-PL potential is trained on PDBBind 2016. It uses a 10 Å cutoff for the pairwise interactions between its 42 ligand, and 23 protein atom types. In the reworked Convex-PL^*R*^ model, we use another version of the original potential, Convex-PL^*cof*^, with a 5.2 Å cutoff, 39 ligand and 33 protein atom types listed in Tables S1, S2 of SI. Truncation of the cutoff radius alone already decreases the virtual screening test bias towards bigger ligands by reducing the variance and the total number of pairwise interactions. We also augmented the training decoys generation procedure by adding decoys with hydrogen atoms randomly substituted by heavy atoms.

To obtain Convex-PL^*cof*^, we collected a dataset consisting of PDBBind 2019,^66^ Binding MOAD 2020,^67^ and a set of complexes from PDB that contained cofactors or modified residues. The latter were chosen based on the PDBe-KB^68^ annotations, with the exclusion of some potential solvents and buffers provided by VHELIBS. ^69^ We chose to use biological assembly submissions as the representatives of the complexes to be consistent with PDBBind and BindingMOAD, although in some cases of binding at the protein-protein interfaces such an approach can result in wrong binding site configurations. Finally, we filtered out complexes with small molecules having low occupancy, RSCC values, and extreme bond length outliers. RSCC values and geometry outliers were taken from the PDB validation reports. This procedure resulted in 29,763 protein-ligand pairs that were used for training. The dataset creation pipeline is described in more detail in Table S3 of SI and is available at https://github.com/chmnk/pl_binding. Some rare ligand-cofactor atom type pairs, however, are not present in the training set. In such cases, we set these interactions to zero and output a warning.

We then trained Convex-PL^*R*^ on a subset of the PDBBind 2019 general dataset. The subset was chosen to contain complexes with Convex-PL^*cof*^ scores that do not correlate sufficiently with the affinities. To do this, we built a simple regression model and removed complexes with scores at a 0.5 threshold distance from the trend line. We have also excluded complexes satisfying a number of criteria listed in Table S4 of SI. The total size of the regression training subset was equal to 12,019 complexes.

## Model assessment and discussion

### CASF Benchmarks 2013 and 2016

We firstly assessed the performance of the obtained scoring functions using the two state-of-the art Comparative Assessment of Scoring Functions (CASF) benchmarks, versions 2013^70^ and 2016.^71^ CASF-2013 benchmark consists of 195 complexes, formed by 65 proteins, each binding to 3 different ligands. To assess the abilities of a scoring function, it suggests four tests. The “docking” test aims at the best near-native pose prediction. The “scoring” test evaluates the abilities of a scoring function to predict relative binding affinities by computing the correlation coefficient between the modeled and experimentally obtained binding constants on a set of 195 complexes. In the “ranking” test, for each protein, the most affine ligand should be found among three candidates. In the “screening” test, the goal is to find the true binders of a protein among a large number of “alien” ligands, whose binding affinity to the target protein is unknown.

CASF-2016 benchmark consists of 285 complexes, formed by 57 proteins, each binding to 5 different ligands. The four provided tests are similar to those from the CASF-2013 benchmark except the “ranking” test, where ligand ranking is evaluated by a comparison of average correlation coefficients between experimental and predicted affinities computed for every five complexes of each cluster. CASF-2013 and CASF-2016 contain evaluation results of 20 and 34 scoring functions, respectively.

Figures 2 and 3 summarize the results of Convex-PL^*R*^ performing on these tests. More details can be found in Tables S6-S8 of the SI. Overall, Convex-PL^*R*^ demonstrates good results, although it does not outperform some of the recently released scoring functions.

**Figure 2:**
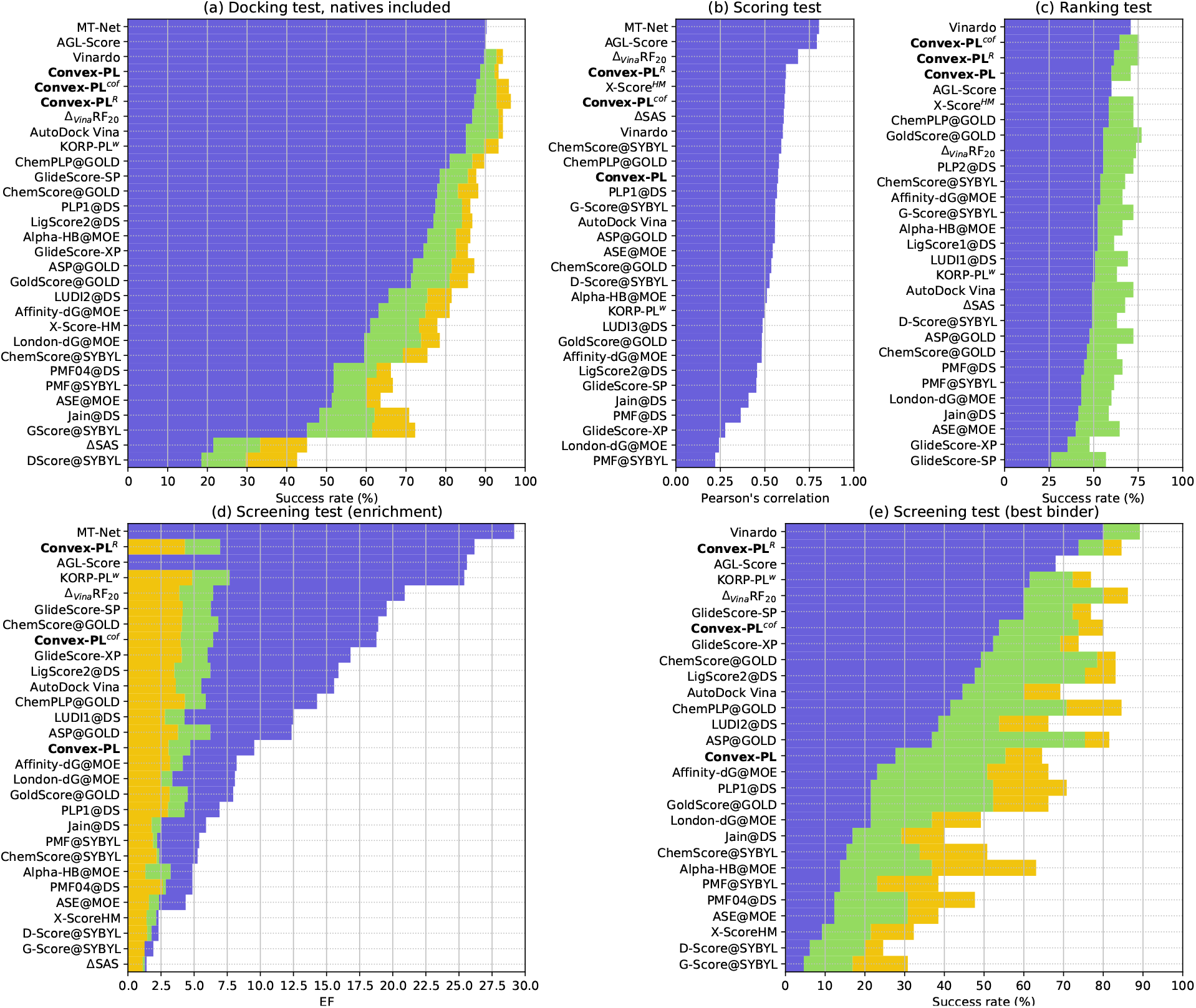
CASF-2013 benchmark results. (a) The success rate of finding a native or near-native pose within 2 Å RMSD in 1 (blue), 2 (green), and 3 (yellow) top-ranked predictions. Pearson’s correlation between predicted scores and experimental log *K*_*a*_ constants. (c) The success rate of the correct ranking of all the three ligands binding the target protein (blue), and ranking the best complex as the top one (green). (d) Enrichment factors computed considering 1% (blue), 5% (green), and 10% (yellow) of the top-ranked compounds. We should note here that Vinardo’s performance was not reported in the corresponding paper^17^. (e) The success rate of identifying the highest-affinity binder among the 1% (blue), 5% (green), or 10% (yellow) top-ranked ligands. We have added our results of KORP-PL^*w*^,^29^ and the evaluations of Vinardo, ^17^ MT-Net,^19^ and AGL-Score, ^25^ found in literature. Not all the test metrics were reported for these scoring functions, resulting in partially empty bars.

**Figure 3:**
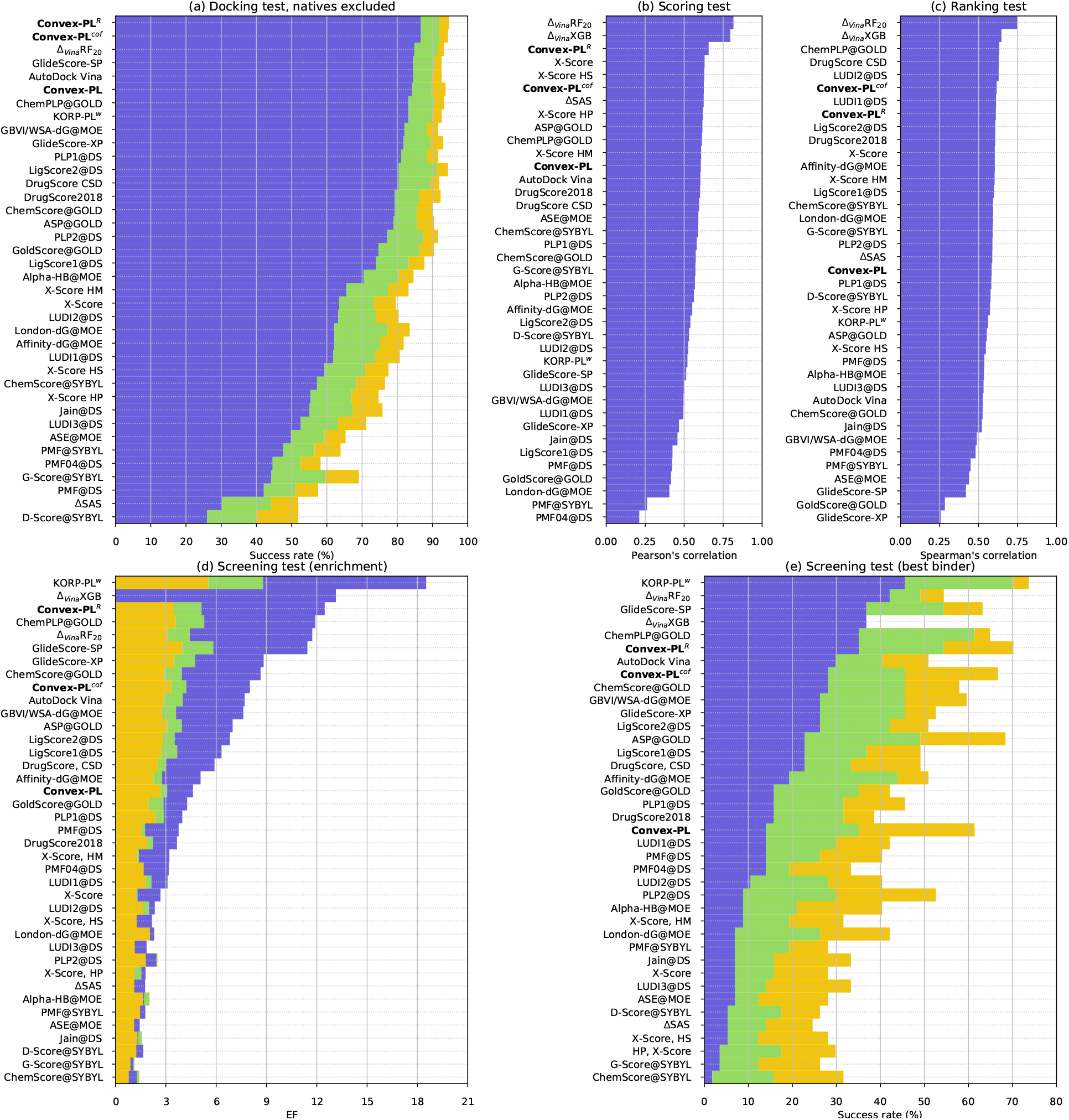
CASF-2016 benchmark results. (a) The success rate of finding a near-native pose within 2 Å RMSD in 1 (blue), 2 (green), and 3 (yellow) top-ranked predictions. Native poses are excluded. (b) Pearson’s correlation with confidence values between predicted scores and experimental log *K*_*a*_. (c) Spearman’s correlation with confidence values computed for scores obtained for every five complexes of the 57 clusters. (d) Enrichment factors computed considering 1% (blue), 5% (green), and 10% (yellow) of the top-ranked compounds. (e) The success rate of identifying the highest-affinity binder among the 1% (blue), 5% (green), or 10% (yellow) top-ranked ligands. We have added our results of KORP-PL^*w*^,^29^ and the evaluations of Δ_*V ina*_XGB,^26^ found in literature. Not all the test metrics were reported for this scoring function, resulting in partially empty bars.

### Scoring and ranking

Convex-PL^*R*^ obtains scores with rather high correlation to the experimental binding constants, which are plotted in Figure 4. Below we discuss several outliers present in the two benchmarks.

**Figure 4:**
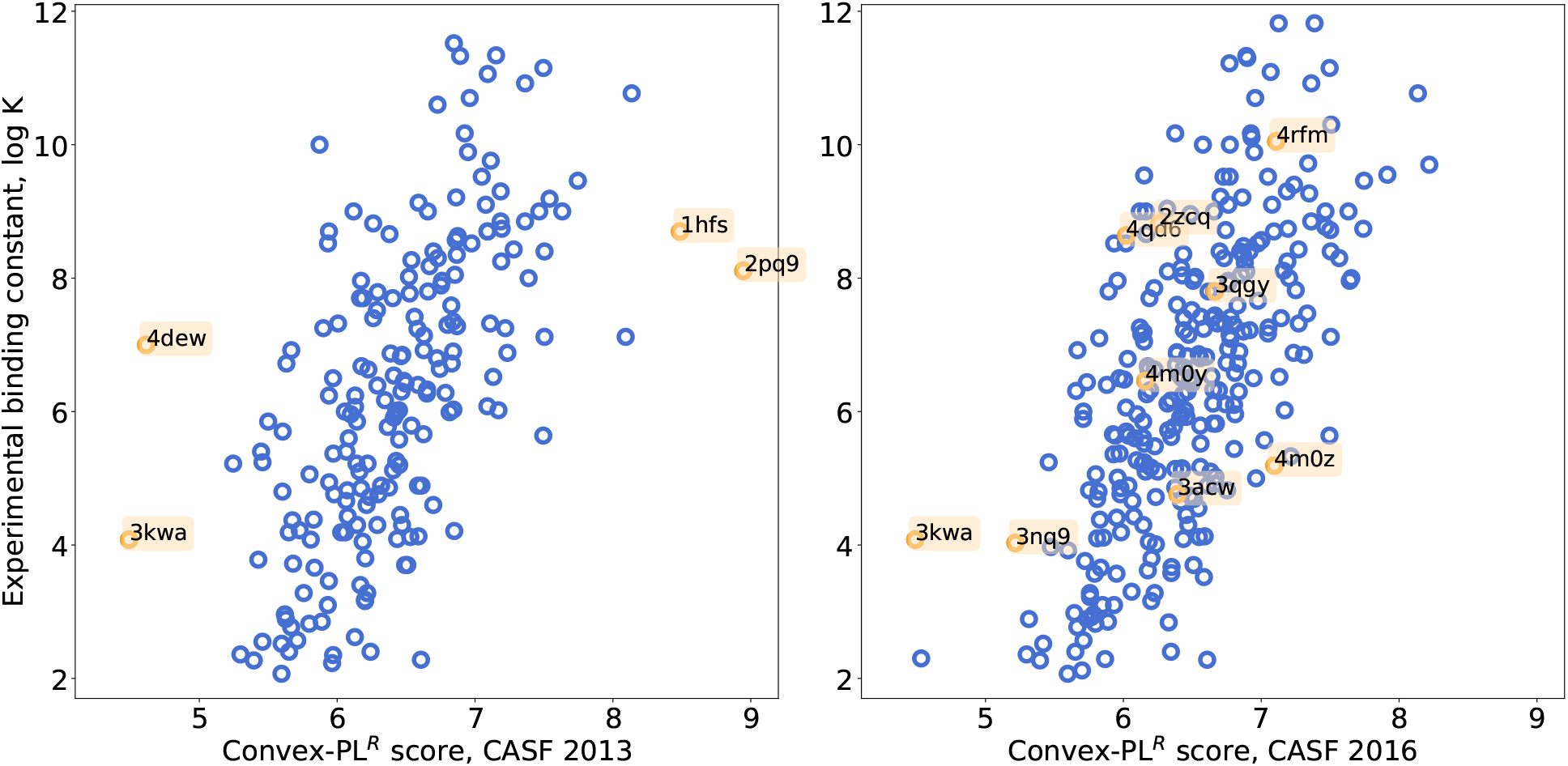
Convex-PL^*R*^ scores, obtained for the CASF benchmarks, versus experimental binding constants. PDB codes of the outliers and challenging complexes discussed in the text are highlighted in orange.

The binding site of the 3kwa complex includes a zinc atom. Although we included some metals into the cofactor-containing dataset, this example may indicate that interactions with zinc are still not fully supported. The binding site of the 4dew complex is located on an interface between the two monomers. This assembly was constructed by applying a crystallographic symmetry operator to the monomeric asymmetric subunit. This resulted in unnaturally small distances (up to 2.5 Å) between the heavy atoms of the ligand aromatic ring and the lysine amino group of the reconstructed monomer, and, in the underestimation of the binding constant. The 1hfs complex consists of a large molecule that binds at the interface of two protein monomers. Similarly to the 4dew example, one of these monomers is not present in the asymmetric unit, so the excessively favourable score produced by Convex-PL^*R*^ can be explained by a possibility of an artefact of the binding interface reconstruction. In the case of 2pq9, we could not find a good explanation of the particularly high score predicted by Convex-PL^*R*^.

Both CASF-2013 and 2016 ranking test performance did not improve much upon supplementing Convex-PL with additional descriptors and even dropped in some cases. However, seven of fifteen clusters of the CASF-2016 benchmark, in which Convex-PL^*R*^ made predictions with average Spearman correlation coefficients less or equal than 0.3, were on average wrongly predicted by the majority of other scoring functions assessed in the benchmark. Below we would like to discuss the reasons for the poor performance of Convex-PL^*R*^ on two clusters.

The 2zcq complex of the 2zcq cluster contains the magnesium ions interacting with the ligand, and it seems that we are overlooking the electrostatic interactions between the two magnesium ions and a phosphonosulfonate ligand headgroup.^72^ Three other proteins of the cluster (4ea2, 2zcr, and 2zy1) are ranked correctly. Notably, all versions of Convex-PL and other top-performing scoring functions overscore the binding affinity of the 3acw complex, which we can hardly interpret.

An interesting case where both Convex-PL^*R*^ and the top-ranked scoring functions, including Δ_Vina_RF_20_, show near-zero correlation is the 4rfm cluster consisting of the 4rfm, 4qd6, 3qgy, 4m0y, and 4m0z PDB structures corresponding to the interleukin-2 inducible T-cell kinase (ITK) inhibitors. Here, most of the scoring functions, including Convex-PL, disfavor the 4qd6 ligand, although its experimental binding affinity is high and should be ranked second. It seems that the structures of the complexes from the benchmark correspond to those marked as a biological assembly in the Protein Data Bank. However, the biological assembly deposited for 4qd6, along with the published analysis of its interactions with the ligand,^73^ does not include or describe the beta-sheet, which seems to be a part of domain swapping and is present in the original crystallographic structure. Conversely, the protein part that was involved in domain swapping as a beta-sheet in 4qd6 adopts another conformation in 4rfm, 3qgy, 4m0y, and 4m0z. In these complexes, it stops participating in domain swapping and forms a beta-sheet inside the main protein chain. As a result, interactions that contribute to the ligand-binding in 4rfm, 3qgy, and 4m0y are lost in 4qd6, because of the incomplete protein structure, leading to lower affinity predictions. The binding affinity score predicted with Convex-PL^*R*^ for the 4qd6 ligand in a complex with the two protein chains instead of one turned out to be closer to the experimental constant. The original 4m0z and 4m0y structures contain ligands that bind to the two sites of ITK,^74^ one of which was chosen per each protein for the benchmark creation. For some reason, Convex-PL^*R*^ and other scoring functions overestimate the affinity of the ligand binding to the allosteric site of the 4m0z structure. We have also noticed that the binding affinity provided in the benchmark for 4m0y corresponds to the allosteric pocket binding, ^74^ while the ligand chosen for the benchmark binds the ATP pocket. The correct choice of the 4m0y affinity also increases the ranking and scoring test performance of Convex-PL^*R*^, as well as of several other scoring functions. This case particularly illustrates the vulnerability of binding energy prediction models to the interpretation of data in various steps of data collection and processing. These include uncertain information in the papers with multiple binding constants coming from different experiments, unreliable biological assembly information, incorrect interpretation of electron density of small molecules, etc.

### Docking and screening tests

Convex-PL performance on the docking test of the CASF-2013 benchmark was discussed in detail before.^16^ Convex-PL^*R*^ keeps and improves its high performance in both CASF benchmarks.

CASF benchmarks suggest an evaluation of the virtual screening performance in terms of enrichment factors. They show how better is the scoring function-based prediction of the true binders compared to random picking and can be defined as follows:

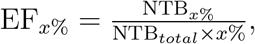

where NTB_*x*%_ is the number of true binders found in the top-x% of the configurations with the highest scores. NTB_*total*_ refers to the total number of true binders for a protein. Improvement of the screening power was the central goal of modifications we applied to Convex-PL, and on these tests, we can clearly see the importance of model re-training and additional descriptors. In the enrichment prediction on CASF-2013, Convex-PL^*R*^ is ranked second after the multi-task MT-NET. In CASF-2016 enrichment prediction, it is outperformed by KORP-PL^*w*^ and Δ_*V ina*_XGB. Overall, the inclusion of additional descriptors and a smaller value of pairwise interactions cutoff value finally allowed us to overcome the bias towards bigger ligands and tighter interfaces, as it is shown in Figure 5.

**Figure 5:**
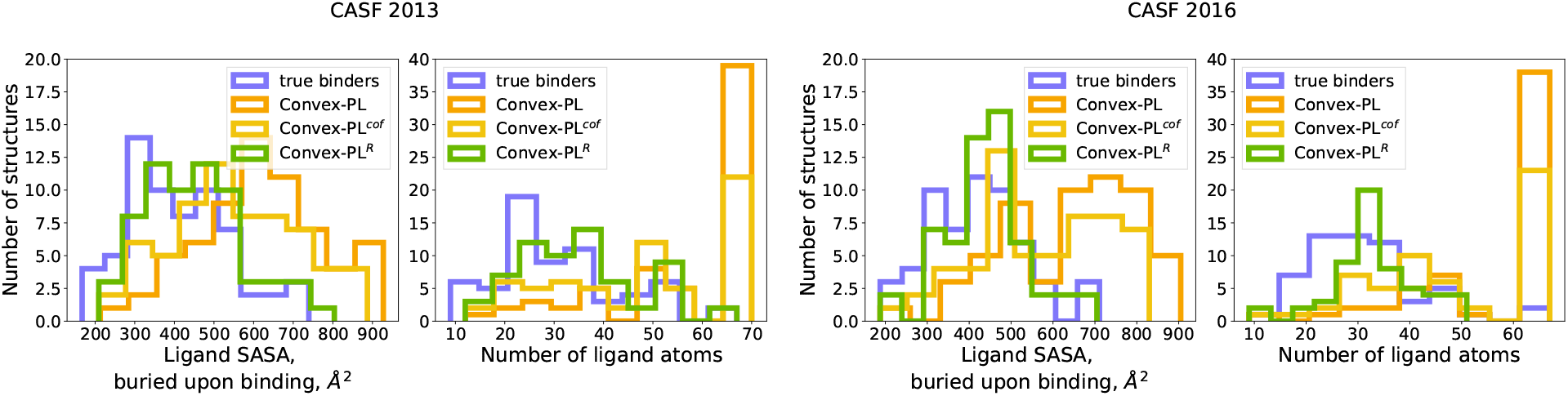
Histograms of average ligand buried SASA and number of atoms computed for the native “truly binding” ligands and decoy poses, top-ranked by different Convex-PL versions in the virtual screening tests from the CASF-2013 and CASF-2016 benchmarks.

### D3R

In addition to the CASF benchmark, we evaluated pose and affinity predictions on a dataset collected from user submissions in the D3R Grand Challenges 2,^75^ 3,^76^ and 4.^77^ The derivation of this test can be found elsewhere. ^29^

Figure 6 demonstrates the high performance of Convex-PL^*R*^ in both pose and affinity prediction parts of the Grand Challenge 2 test. In the Grand Challenge 3 exercise, both Convex-PL^*R*^ and Convex-PL^*cof*^ outperform their predecessor in pose prediction. During the D3R Grand Challenge 3 timeframe, only a few protocols that did not use visual inspection and ligand-based approaches were successful,^76^ meaning that this particular case could be a challenge for many scoring functions. Convex-PL tended to prefer incorrect poses that were buried deeper in the binding pocket and had tighter binding interfaces. Convex-PL^*R*^ and Convex-PL^*cof*^ partially overcome this problem. However, their ability to top-score the correct near-native pose is still lower than that of KORP-PL^*w*^. In the affinity prediction exercise of Grand Challenge 3, additional penalization introduced in Convex-PL^*R*^ seems to worsen the correlations in comparison with Convex-PL^*cof*^. Finally, in Grand Challenge 4 the model without entropic contributions and original Convex-PL turned out to perform better in pose prediction.

**Figure 6:**
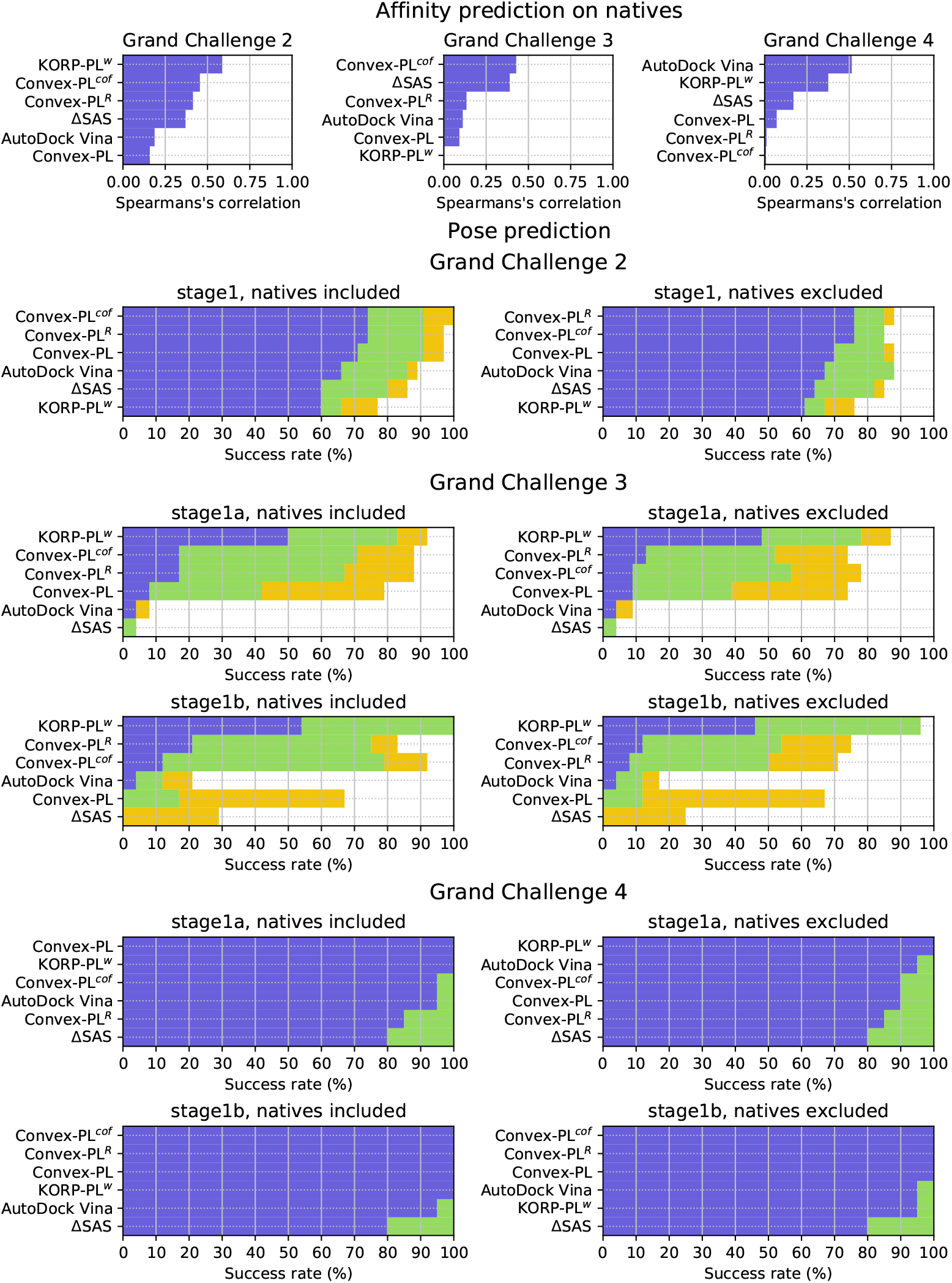
D3R pose prediction and scoring results. Success rates of finding a pose within 2 Å RMSD from the native conformation among the 1%, 5%, and 10% of top-ranked poses are shown in blue, green, and yellow, respectively. Scoring power is represented by the Spearman’s correlation coefficient between the predicted and experimental binding constants. These success rates are computed with respect to the actual number of ligands, for which the poses with the desired RMSD values were present in the user submissions. More detailed results can be found in Tables S9-S14 of SI.

### DUD

To additionally measure the virtual screening performance of Convex-PL^*R*^, we assessed it using the Directory of Useful Decoys (DUD)^78^ dataset specifically designed for virtual screening benchmarking. It contains 40 targets with a total of 2,950 active compounds. For each active compound, the dataset provides 36 decoys with similar physics, but various chemical topology. In this test, we assessed Convex-PL^*R*^, Convex-PL^*cof*^, and Convex-PL scoring functions together with a few other ones, namely, KORP-PL^*w*^, AutoDock Vina, and Smina^79^ with Vinardo chosen as the scoring function.

### Docking protocol and results

For each of the DUD targets, we ran docking simulations using *VinaCPL*, an in-house modified version of AutoDock Vina with the original Convex-PL potential as a scoring function. Since Convex-PL is pairwise and distance-dependent, it could be naturally mapped to the AutoDock Vina pairwise interaction grids. In more detail, we updated Vina’s atom types to the 42 ligand and 23 protein types and replaced the AutoDock Vina scoring function with Convex-PL. We additionally penalized intra-ligand clashes using the values of the Convex-PL protein-ligand potential mapped to the ligand-ligand atom types. Our modifications of AutoDock Vina also increased the *num saved min* internal parameter and skipped the RMSD-based clustering, so that more conformations could be generated for further re-scoring dependent on the provided *num modes* command-line argument.

We then re-scored the obtained docking poses with Convex-PL^*R*^, Convex-PL^*cof*^, and KORP-PL^*w*^. We also performed the rescoring with the original Convex-PL to obtain the scores that do not include intra-ligand clashes. The final score of each compound was obtained by averaging the scores of the top-10 Convex-PL^*R*^ predictions. We have already successfully applied earlier versions of this protocol in the D3R Grand Challenges 2^80^ and 4.^81^

AutoDock Vina, Smina, and VinaCPL were launched with the exhaustiveness parameter set to 10, other parameters except for the number of output conformations in the case of VinaCPL were left to their defaults. We have tested several ways to define the docking binding boxes. We achieved the best virtual screening results for all three protocols with the binding box determined with the following procedure: *(i)* Target co-crystal ligand dimensions, namely the co-crystal box, were measured. *(ii)* Dimensions of the input ligand 3D conformation provided in the dataset, namely the ligand box, were measured. *(iii)* The input ligand box was aligned with the co-crystal box. *(iv)* Finally, for each dimension, the maximum size of the two boxes was chosen and multiplied by an arbitrarily chosen factor of Notably, virtual screening results of both Vina and Vinardo achieved with such a box outperformed those reported previously.^17^

Figure 7, Table 1, and the Tables S16-S17 of SI summarize virtual screening powers evaluated by measuring the ROC AUC, enrichment factors, and BEDROC_*α*=20_. Here, both Convex-PL^*R*^ and Convex-PL^*cof*^ considerably improve the Convex-PL performance and out-perform both Vina and Vinardo in early enrichment metrics. However, their results are still worse than those of KORP-PL^*w*^. In more discriminative enrichment metrics, Convex-PL^*R*^, with its entropic terms, is surprisingly outperformed by Convex-PL^*cof*^. On DUD, where decoy molecules were specifically generated and do not have such considerable size difference from actives as in some CASF targets, the benefit introduced by an entropic term can have a smaller contribution. In five targets that contain cofactors, both new versions of Convex-PL outperform KORP-PL^*w*^, which does not have cofactor support. However, in four other cofactor-containing targets, Convex-PL^*R*^ and Convex-PL^*cof*^ demonstrated rather low results. This could be explained by the fact that VinaCPL, which we used for docking, does not have the cofactor support yet.

**Figure 7:**
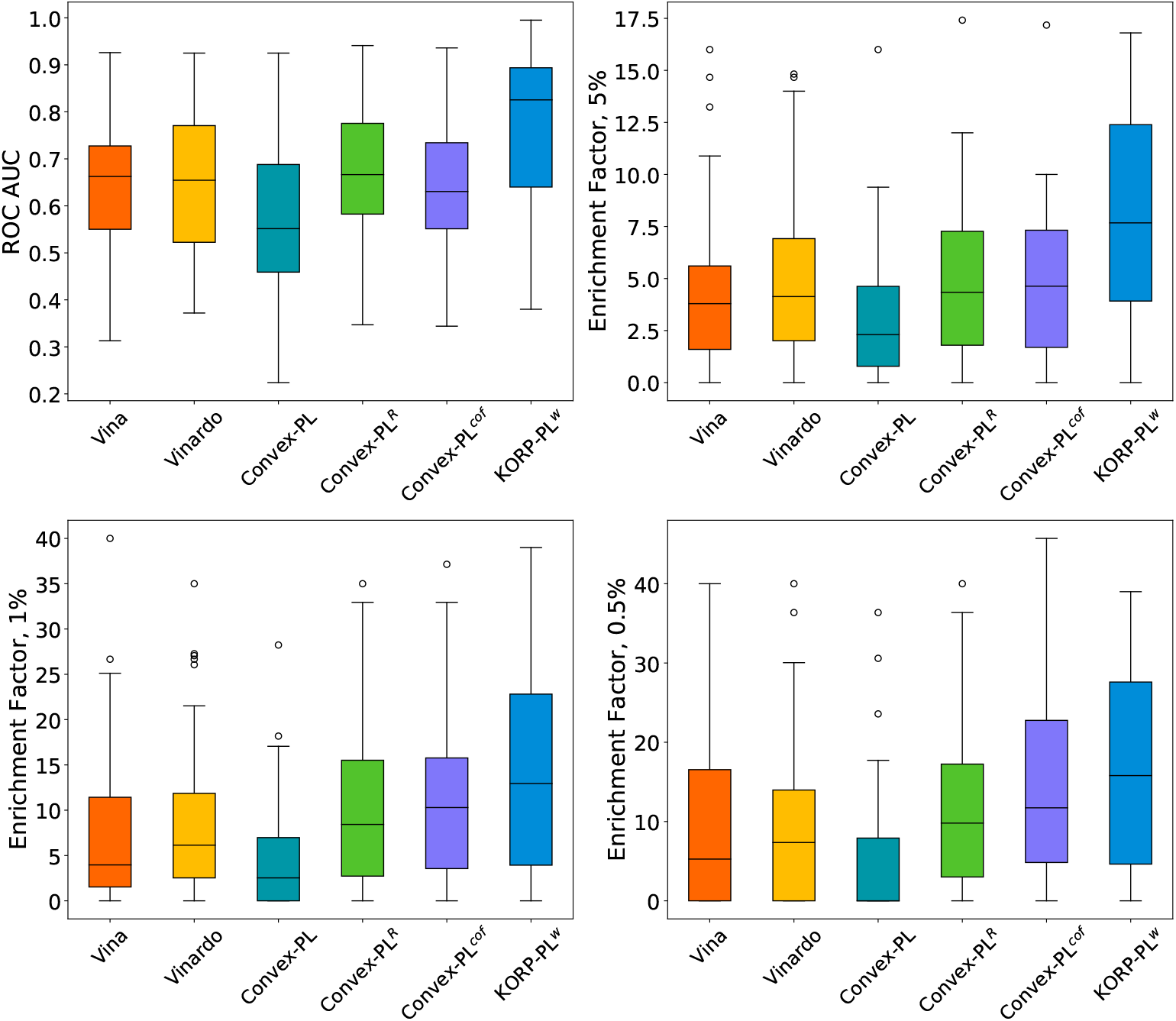
Box plots representing the ROC AUC, and 5%, 1%, 0.5% enrichment factors computed for the DUD dataset with AutoDock Vina, Vinardo, Convex-PL^*R*^, Convex-PL^*cof*^, Convex-PL, and KORP-PL^*w*^.

**Table 1:**
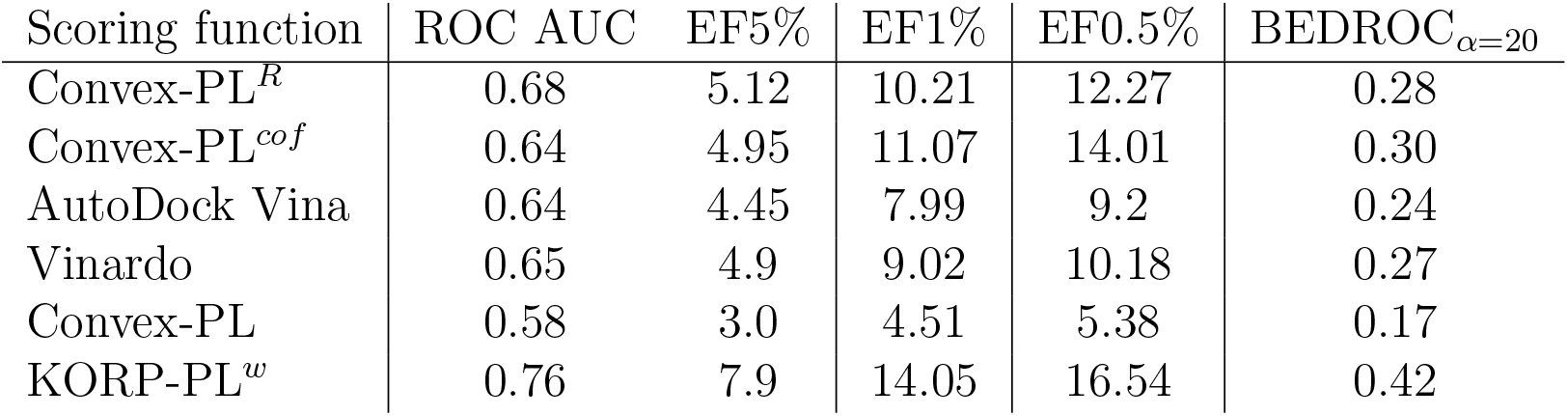
Average ROC AUC scores, 5%, 1%, 0.5% enrichment factor, and BEDROC^82,83^ values computed for the DUD dataset. Per-target evaluation results can be found in Tables S16-S17 of SI. Cofactor-containing targets are listed in Table S15.

### LIT-PCBA

Known drawbacks of DUD include the unproved inactivity of all the decoys and the biases of decoy generation. Therefore, we have also evaluated Convex-PL^*R*^ on the LIT-PCBA virtual screening benchmark^84^ that contains 15 targets with 9,954 active and 2,767,111 inactive compounds found in biological assay databases. Here we compare only AutoDock Vina and Convex-PL^*R*^.

We used the same docking protocols as in the case of DUD. 3D ligand conformers were generated using RDKit’s *EmbedMolecule* with *ETKDGv3* parametrization.^83,85^ Prior to docking, we removed duplicating ligands having different PubChem substance IDs but encoded with the same SMILES strings. We also used only a part of the target representative proteins listed in Table S18, selected from their binding pocket alignment and visual analysis. For each ligand, we took the top-scored predictions among the chosen protein representatives.

Figure 8 and Table 2 demonstrate that Convex-PL^*R*^ outperforms AutoDock Vina in this benchmark. However, their ROC AUC values and enrichment factors are, in general, rather low. Docking to 7 and 5 targets for Vina and Convex-PL^*R*^, respectively, resulted in zero 0.5% enrichment. We suppose that such poor results can be partially related to the benchmark composition. For example, the *kat2a* target contains very different representatives with binding pockets located on two different domains, while the *adrb2* representatives contain proteins that were specifically mutated for covalent binding. Similarly, the *aldh1, idh1*, and *mtorc1* targets have two binding pockets, *vdr* has a mutation in a binding site. Although, in principle, labeled inactive compounds are much more valuable than decoys, it is not clear whether all of the hundred thousands of actives and inactives were properly validated against the particular targets, as a considerable portion of the benchmark is based on cell-based assays.

**Figure 8:**
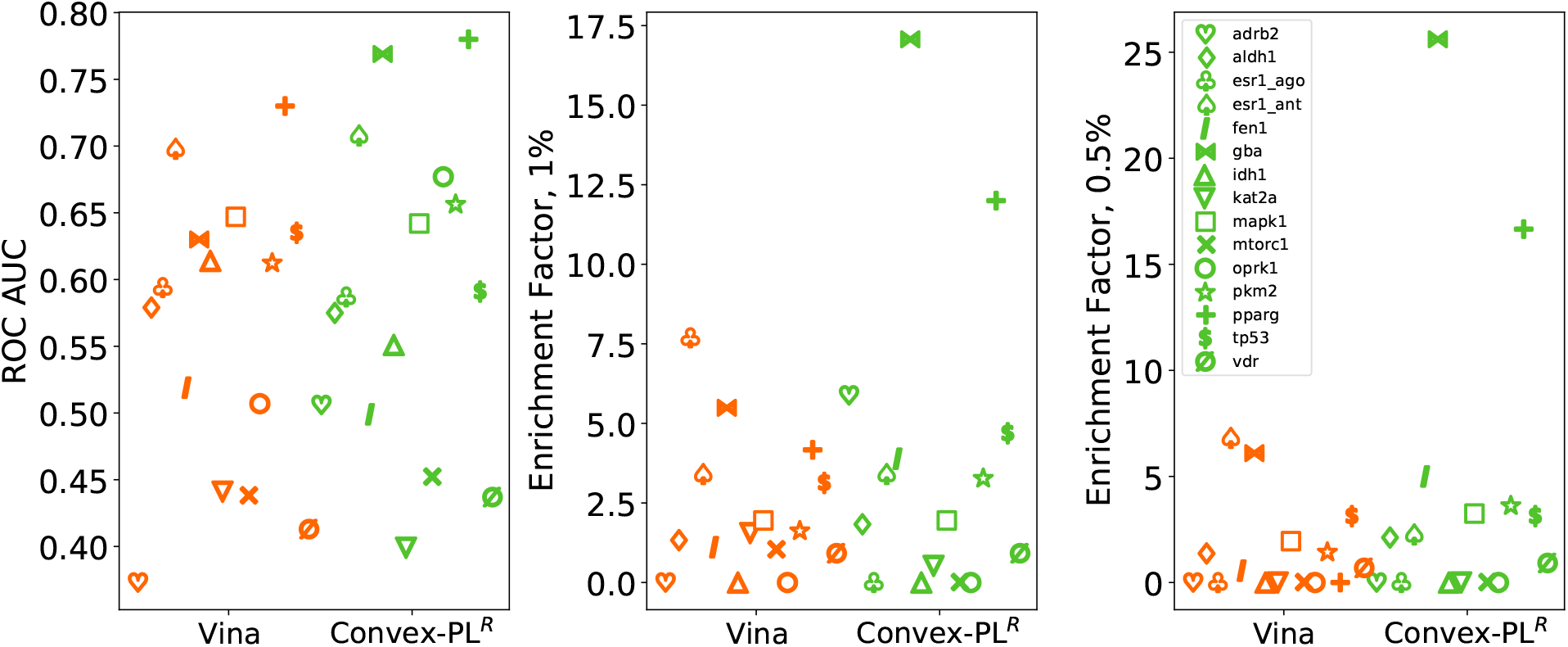
ROC AUC, and 1%, 0.5% enrichment factors computed for the LIT-PCBA dataset with AutoDock Vina, and Convex-PL^*R*^.

**Table 2:** Average ROC AUC scores, 5%, 1%, 0.5% enrichment factor, and BEDROC^82,83^ values computed for the LIT-PCBA dataset. Per-target evaluation results can be found in Table S19 of SI.

| Scoring function | ROC AUC | EF5% | EF1% | EF0.5% | BEDROC<sub>$\alpha=20$</sub> |
| --- | --- | --- | --- | --- | --- |
| AutoDock Vina | 0.562 | 0.086 | 1.572 | 2.23 | 1.47 |
| Convex-PL<sup>R</sup> | 0.6 | 0.111 | 2.159 | 3.70 | 4.18 |

### Technical details

Convex-PL^*R*^ is written in C++ and is available as a standalone binary. It takes about 16 milliseconds on a single core of Linux Intel(R) Core(TM) i7-8565U CPU @ 1.80GHz to score a protein-ligand complex from the CASF-2013 core set containing a single ligand pose of 25 heavy atoms on average. Convex-PL^*cof*^ is about 1.5 times faster, as it does not require the SASA computation.

## Conclusion

This paper presents Convex-PL^*R*^ – a reworked Convex-PL protein-ligand potential, derived from thermodynamical considerations. Our model incorporates conformational entropic terms for the ligand and the binding pocket sidechain flexibility. For the solvation contributions, we have developed two models, either computing atom-mesh interactions, or approximating the solvation energy with SASA. To our surprise, the weights of these contributions turned out to be negligibly small after the training, and we removed them from the final model. We also developed a novel docking protocol by incorporating the Convex-PL scoring function inside Autodock Vina with a subsequent rescoring of the docking poses with Convex-PL^*R*^. We successfully validated Convex-PL^*R*^ on CASF 2013 and 2016 benchmarks, on a dataset derived from the D3R Challenges, and on DUD and LIT-PCBA virtual screening tests, where it was ranked top-2 – top-3 in the majority of the CASF tests, top-2 in DUD, D3R, and outperformed AutoDock Vina in LIT-PCBA.

## Abbreviations

CASF: Comparative Assessment of Scoring Functions
RMSD: Root Mean Squared Deviation
SVM: Support Vector Machinesn
PDB: Protein Data Bankn
SAS: Solvent-Accessible Surfacen
SASA: Solvent-Accessible Surface Area

## Acknowledgement

The authors would like to thank Ivan Gushchin from MIPT Moscow for providing his expertise in crystallography, Pablo Pablo Chacón from Rocasolano Institute of Physical Chemistry, Madrid for the discussions of the representation of the desolvation free energy change in knowledge-based potentials, and anonymous reviewers for their comments and evaluation. V.Ch. is supported by the Ministry of Science and Higher Education of the Russian Federation (agreement # 075-00337-20-03, project FSMG-2020-0003).

## References

(1) Böhm, H.-J. (1994) The Development of a Simple Empirical Scoring Function to Estimate the Binding Constant for a Protein-Ligand Complex of Known Three-Dimensional Structure. J. Comput.-Aided Mol. Des. 8, 243–256.

(2) Jones, G., Willett, P., Glen, R. C., Leach, A. R., and Taylor, R. (1997) Development and validation of a genetic algorithm for flexible docking. J. Mol. Biol. 267, 727–748.

(3) Eldridge, M. D., Murray, C. W., Auton, T. R., Paolini, G. V., and Mee, R. P. (1997) Empirical Scoring Functions: I. The Development of a Fast Empirical Scoring Function to Estimate the Binding Affinity of Ligands in Receptor Complexes. J. Comput.-Aided Mol. Des. 11, 425–445.

(4) Gohlke, H., Hendlich, M., and Klebe, G. (2000) Knowledge-Based Scoring Function to Predict Protein-Ligand Interactions. J. Mol. Biol. 295, 337–356.

(5) Jain, A. N. (1996) Scoring noncovalent protein-ligand interactions: a continuous differentiable function tuned to compute binding affinities. J. Comput.-Aided Mol. Des. 10, 427–440.

(6) Wang, R., Lai, L., and Wang, S. (2002) Further Development and Validation of Empirical Scoring Functions for Structure-Based Binding Affinity Prediction. J. Comput.- Aided Mol. Des. 16, 11–26.

(7) Friesner, R. A., Murphy, R. B., Repasky, M. P., Frye, L. L., Greenwood, J. R., Halgren, T. A., Sanschagrin, P. C., and Mainz, D. T. (2006) Extra precision glide: Docking and scoring incorporating a model of hydrophobic enclosure for proteinligand complexes. J. Med. Chem. 49, 6177–6196.

(8) Kulharia, M., Goody, R. S., and Jackson, R. M. (2008) Information theory-based scoring function for the structure-based prediction of protein-ligand binding affinity. J. Chem. Inf. Model. 48, 1990–1998.

(9) Trott, O., and Olson, A. J. (2010) AutoDock Vina: Improving the Speed and Accuracy of Docking with a New Scoring Function, Efficient Optimization, and Multithreading. J. Comput. Chem. 31, 455–461.

(10) Ballester, P. J., and Mitchell, J. B. (2010) A machine learning approach to predicting protein-ligand binding affinity with applications to molecular docking. Bioinformatics 26, 1169–1175.

(11) Korb, O., Stutzle, T., and Exner, T. E. (2009) Empirical scoring functions for advanced protein–ligand docking with PLANTS. J. Chem. Inf. Model. 49, 84–96.

(12) Neudert, G., and Klebe, G. (2011) DSX: a knowledge-based scoring function for the assessment of protein–ligand complexes. J. Chem. Inf. Model. 51, 2731–2745.

(13) Verdonk, M. L., Ludlow, R. F., Giangreco, I., and Rathi, P. C. (2016) Protein-ligand in-formatics force field (PLIff): Toward a fully knowledge driven “force field” for biomolecular interactions. J. Med. Chem. 59, 6891–6902.

(14) Zilian, D., and Sotriffer, C. A. (2013) SFCscore RF: a random forest-based scoring function for improved affinity prediction of protein–ligand complexes. J. Chem. Inf. Model 53, 1923–1933.

(15) Debroise, T., Shakhnovich, E. I., and Cheron, N. (2017) A hybrid knowledge-based and empirical scoring function for protein–ligand interaction: SMoG2016. J. Chem. Inf. Model 57, 584–593.

(16) Kadukova, M., and Grudinin, S. (2017) Convex-PL: a novel knowledge-based potential for protein-ligand interactions deduced from structural databases using convex optimization. J. Comput.-Aided Mol. Des. 31, 943–958.

(17) Quiroga, R., and Villarreal, M. A. (2016) Vinardo: A Scoring Function Based on Autodock Vina Improves Scoring, Docking, and Virtual Screening. PloS One 11, e0155183.

(18) Wallach, I., Dzamba, M., and Heifets, A. (2015) AtomNet: a deep convolutional neural network for bioactivity prediction in structure-based drug discovery. arXiv preprint 1510.02855

(19) Ashtawy, H. M., and Mahapatra, N. R. (2017) Task-Specific Scoring Functions for Predicting Ligand Binding Poses and Affinity and for Screening Enrichment. J. Chem. Inf. Model 58, 119–133.

(20) Li, G.-B., Yang, L.-L., Wang, W.-J., Li, L.-L., and Yang, S.-Y. (2013) ID-Score: A New Empirical Scoring Function Based on a Comprehensive Set of Descriptors Related to Protein–Ligand Interactions. J. Chem. Inf. Model. 53, 592–600, PMID: 23394072.

(21) Wang, S.-H., Wu, Y.-T., Kuo, S.-C., and Yu, J. (2013) HotLig: A molecular surface-directed approach to scoring protein–ligand interactions. J. Chem. Inf. Model. 53, 181–2195.

(22) Wang, C., and Zhang, Y. (2017) Improving scoring-docking-screening powers of protein– ligand scoring functions using random forest. J. Comput. Chem. 38, 169–177.

(23) Jim’senez, J., Skalic, M., Martinez-Rosell, G., and De Fabritiis, G. (2018) KDEEP: Protein–ligand absolute binding affinity prediction via 3D-convolutional neural networks. J. Chem. Inf. Model. 58, 287–296.

(24) Yan, Z., and Wang, J. (2016) Incorporating specificity into optimization: evaluation of SPA using CSAR 2014 and CASF 2013 benchmarks. J. Comput.-Aided Mol. Des. 30, 219–227.

(25) Nguyen, D. D., and Wei, G.-W. (2019) Agl-score: Algebraic graph learning score for protein–ligand binding scoring, ranking, docking, and screening. J. Chem. Inf. Model 59, 3291–3304.

(26) Lu, J., Hou, X., Wang, C., and Zhang, Y. (2019) Incorporating Explicit Water Molecules and Ligand Conformation Stability in Machine-Learning Scoring Functions. J. Chem. Inf. Model 59, 4540–4549.

(27) Chen, P., Ke, Y., Lu, Y., Du, Y., Li, J., Yan, H., Zhao, H., Zhou, Y., and Yang, Y. (2019) DLIGAND2: an improved knowledge-based energy function for protein–ligand interactions using the distance-scaled, finite, ideal-gas reference state. J. Cheminformatics 11, 52.

(28) Karlov, D. S., Sosnin, S., Fedorov, M. V., and Popov, P. (2020) graphDelta: MPNN Scoring Function for the Affinity Prediction of Protein–Ligand Complexes. ACS Omega 5, 5150–5159.

(29) Kadukova, M., Machado, K. d. S., Chacón, P., and Grudinin, S. (2020) KORP-PL: a coarse-grained knowledge-based scoring function for protein-ligand interactions. Bioinformatics In press.

(30) Francoeur, P., Masuda, T., Sunseri, J., Jia, A., Iovanisci, R. B., Snyder, I., and Koes, D. R. (2020) Three-Dimensional Convolutional Neural Networks and a Cross-Docked Data Set for Structure-Based Drug Design. J. Chem. Inf. Model 60, 4200–4215.

(31) Scantlebury, J., Brown, N., Von Delft, F., and Deane, C. M. (2020) Dataset Augmentation Allows Deep Learning-Based Virtual Screening To Better Generalize To Unseen Target Classes, And Highlight Important Binding Interactions. J. Chem. Inf. Model 60, 3722–3730.

(32) Gilson, M. K., and Zhou, H.-X. (2007) Calculation of protein-ligand binding affinities. Annu. Rev. Biophys. Biomol. Struct. 36, 21–42.

(33) Gilson, M. K., Given, J. A., Bush, B. L., and McCammon, J. A. (1997) The statistical-thermodynamic basis for computation of binding affinities: a critical review. Biophys. J. 72, 1047–1069.

(34) Li, Y., Su, M., Liu, Z., Li, J., Liu, J., Han, L., and Wang, R. (2018) Assessing protein– ligand interaction scoring functions with the CASF-2013 benchmark. Nat. Protoc. 13, 666.

(35) Su, M., Yang, Q., Du, Y., Feng, G., Liu, Z., Li, Y., and Wang, R. (2018) Comparative assessment of scoring functions: the CASF-2016 update. J. Chem. Inf. Model 59, 895–913.

(36) Schrödinger, LLC, The PyMOL Molecular Graphics System, Version 2.4. 2020.

(37) Ruiz-Carmona, S., Alvarez-Garcia, D., Foloppe, N., Garmendia-Doval, A. B., Juhos, S., Schmidtke, P., Barril, X., Hubbard, R. E., and Morley, S. D. (2014) rDock: A Fast, Versatile and Open Source Program for Docking Ligands to Proteins and Nucleic Acids. PLoS Comput. Biol. 10, 1–7.

(38) Huang, S.-Y., and Zou, X. (2010) Inclusion of Solvation and Entropy in the Knowledge– Based Scoring Function for Protein–Ligand Interactions. J. Chem. Inf. Model. 50, 262–273, PMID: 20088605.

(39) Guedes, I. A., Barreto, A. M., Marinho, D., Krempser, E., Kuenemann, M. A., Sperandio, O., Dardenne, L. E., and Miteva, M. A. (2021) New machine learning and physics-based scoring functions for drug discovery. Sci. Rep. 11, 1–19.

(40) Abagyan, R., and Totrov, M. (1994) Biased probability Monte Carlo conformational searches and electrostatic calculations for peptides and proteins. J. Mol. Biol. 235, 983–1002.

(41) Krissinel, E., and Henrick, K. Detection of protein assemblies in crystals. International Symposium on Computational Life Science. 2005; pp 163–174.

(42) Murray, C. W., and Verdonk, M. L. (2002) The consequences of translational and rotational entropy lost by small molecules on binding to proteins. J. Comput.-Aided Mol. Des. 16, 741–753.

(43) Wang, J., Wang, W., Huo, S., Lee, M., and Kollman, P. A. (2002) Solvation model based on weighted solvent accessible surface area. J. Phys. Chem. B 105, 5055–5067.

(44) Ben-Naim, A. (1997) Statistical Potentials Extracted from Protein Structures: Are These Meaningful Potentials? J. Chem. Phys. 107, 3698–3706.

(45) Lazaridis, T., and Karplus, M. (2003) Effective energy function for proteins in solution. Proteins Struct. Funct. Genet. 52, 176–192.

(46) Labute, P. (2008) The generalized Born/volume integral implicit solvent model: estimation of the free energy of hydration using London dispersion instead of atomic surface area. J. Comput. Chem. 29, 1693–1698.

(47) Yin, S., Biedermannova, L., Vondrasek, J., and Dokholyan, N. V. (2008) MedusaScore: An accurate force field-based scoring function for virtual drug screening. J. Chem. Inf. Model. 48, 1656–1662.

(48) Reulecke, I., Lange, G., Albrecht, J., Klein, R., and Rarey, M. (2008) Towards an integrated description of hydrogen bonding and dehydration: decreasing false positives in virtual screening with the HYDE scoring function. ChemMedChem 3, 885–897.

(49) Schneider, N., Lange, G., Hindle, S., Klein, R., and Rarey, M. (2013) A consistent description of HYdrogen bond and DEhydration energies in protein–ligand complexes: methods behind the HYDE scoring function. J. Comput.-Aided Mol. Des. 27, 15–29.

(50) Cao, Y., and Li, L. (2014) Improved protein–ligand binding affinity prediction by using a curvature-dependent surface-area model. Bioinformatics 30, 1674–1680.

(51) Genheden, S., Luchko, T., Gusarov, S., Kovalenko, A., and Ryde, U. (2010) An MM/3D-RISM approach for ligand binding affinities. J. Phys. Chem. B 114, 8505–8516.

(52) Swanson, J. M., Henchman, R. H., and McCammon, J. A. (2004) Revisiting Free Energy Calculations: A Theoretical Connection to MM/PBSA and Direct Calculation of the Association Free Energy. Biophys. J. 86, 67–74.

(53) Zou, X., Sun, Y., and Kuntz, I. D. (1999) Inclusion of solvation in ligand binding free energy calculations using the generalized-born model. J. Am. Chem. Soc. 121, 8033–8043.

(54) Lee, M. S., and Olson, M. A. (2006) Calculation of absolute protein-ligand binding affinity using path and endpoint approaches. Biophys. J. 90, 864–877.

(55) Totrov, M., and Abagyan, R. (2001) Rapid boundary element solvation electrostatics calculations in folding simulations: Successful folding of a 23-residue peptide. Biopolym. - Pept. Sci. Sect. 60, 124–133.

(56) Schrödinger Release 2019-1: WaterMap. https://www.schrodinger.com/watermap.

(57) Cappel, D., Sherman, W., and Beuming, T. (2017) Calculating Water Thermodynamics in the Binding Site of Proteins–Applications of WaterMap to Drug Discovery. Curr. Top. Med. Chem 17, 2586–2598.

(58) FLAP/WaterFLAP. http://www.moldiscovery.com/software/flap/.

(59) Sridhar, A., Ross, G. A., and Biggin, P. C. (2017) Waterdock 2.0: Water placement prediction for Holo-structures with a pymol plugin. PloS One 12, e0172743.

(60) Lensink, M. F. et al. (2014) Blind prediction of interfacial water positions in CAPRI. Proteins: Struct., Funct., Bioinf. 82, 620–632.

(61) Li, Y., Gao, Y. D., Holloway, M. K., and Wang, R. (2020) Prediction of the Favorable Hydration Sites in a Protein Binding Pocket and Its Application to Scoring Function Formulation. J. Chem. Inf. Model 60, 4359–4375.

(62) Giordanetto, F., Cotesta, S., Catana, C., Trosset, J.-Y., Vulpetti, A., Stouten, P. F., and Kroemer, R. T. (2004) Novel scoring functions comprising QXP, SASA, and protein side-chain entropy terms. J. Chem. Inf. Comput. Sci. 44, 882–893.

(63) Sternberg, M. J., and Chickos, J. S. (1994) Protien side-chain conformational entropy derived from fusion data-comparison with other empirical scales. Protein Eng. Des. Sel. 7, 149–155.

(64) Klenin, K. V., Tristram, F., Strunk, T., and Wenzel, W. (2011) Derivatives of molecular surface area and volume: Simple and exact analytical formulas. J. Comput. Chem. 32, 2647–2653.

(65) Klenin, K., Tristram, F., Strunk, T., and Wenzel, W. (2012) Achieving Numerical Stability in Analytical Computation of the Molecular Surface and Volume. From Computational Biophysics to Systems Biology (CBSB11)–Celebrating Harold Scheraga’s 90th Birthday 8, 75.

(66) Liu, Z., Su, M., Han, L., Liu, J., Yang, Q., Li, Y., and Wang, R. (2017) Forging the basis for developing protein–ligand interaction scoring functions. Acc. Chem. Res. 50, 302–309.

(67) Benson, M. L., Smith, R. D., Khazanov, N. A., Dimcheff, B., Beaver, J., Dresslar, P., Nerothin, J., and Carlson, H. A. (2007) Binding MOAD, a high-quality protein–ligand database. Nucleic Acids Res. 36, D674–D678.

(68) consortium, P.-K. (2019) PDBe-KB: a community-driven resource for structural and functional annotations. Nucleic Acids Research 48, D344–D353.

(69) Cereto-Massagu’se, A., Ojeda, M. J., Joosten, R. P., Valls, C., Mulero, M., Salvado, M. J., Arola-Arnal, A., Arola, L., Garcia-Vallvé, S., and Pujadas, G. (2013) The good, the bad and the dubious: VHELIBS, a validation helper for ligands and binding sites. J. Cheminform. 5, 1–9.

(70) Li, Y., Han, L., Liu, Z., and Wang, R. (2014) Comparative Assessment of Scoring Functions on an Updated Benchmark: 2. Evaluation Methods an General Results. J. Chem. Inf. Model. 54, 1717–36.

(71) Su, M., Yang, Q., Du, Y., Feng, G., Liu, Z., Li, Y., and Wang, R. (2018) Comparative assessment of scoring functions: the CASF-2016 update. J. Chem. Inf. Model 59, 895–913.

(72) Liu, C.-I., Liu, G. Y., Song, Y., Yin, F., Hensler, M. E., Jeng, W.-Y., Nizet, V., Wang, A. H.-J., and Oldfield, E. (2008) A cholesterol biosynthesis inhibitor blocks Staphylococcus aureus virulence. Science 319, 1391–1394.

(73) Trani, G. et al. (2014) Design, synthesis and structure–activity relationships of a novel class of sulfonylpyridine inhibitors of Interleukin-2 inducible T-cell kinase (ITK). Bioorg. Med. Chem. Lett 24, 5818–5823.

(74) Han, S. et al. (2014) Selectively targeting an inactive conformation of interleukin-2-inducible T-cell kinase by allosteric inhibitors. Biochem. J 460, 211–222.

(75) Gaieb, Z., Liu, S., Gathiaka, S., Chiu, M., Yang, H., Shao, C., Feher, V. A., Walters, W. P., Kuhn, B., Rudolph, M. G., Burley, S. K., Gilson, M. K., and Amaro, R. E. (2018) D3R Grand Challenge 2: blind prediction of protein–ligand poses, affinity rankings, and relative binding free energies. J. Comput.-Aided Mol. Des. 32, 1–20.

(76) Gaieb, Z., Parks, C. D., Chiu, M., Yang, H., Shao, C., Walters, W. P., Lambert, M. H., Nevins, N., Bembenek, S. D., Ameriks, M. K., Mirzadegan, T., Burley, S. K., Amaro, R. E., and Gilson, M. K. (2019) D3R Grand Challenge 3: blind prediction of protein–ligand poses and affinity rankings. J. Comput.-Aided Mol. Des. 33, 1–18.

(77) Parks, C. D., Gaieb, Z., Chiu, M., Yang, H., Shao, C., Walters, W. P., Jansen, J. M., McGaughey, G., Lewis, R. A., Bembenek, S. D., Ameriks, M. K., Mirzadegan, T., Burley, S. K., Amaro, R. E., and Gilson, M. K. (2020) D3R Grand Challenge 4: blind prediction of protein–ligand poses, affinity rankings, and relative binding free energies. J. Comput.-Aided Mol. Des.

(78) Huang, N., Shoichet, B. K., and Irwin, J. J. (2006) Benchmarking sets for molecular docking. J. Med. Chem. 49, 6789–6801.

(79) Koes, D. R., Baumgartner, M. P., and Camacho, C. J. (2013) Lessons learned in empirical scoring with smina from the CSAR 2011 benchmarking exercise. J. Chem. Inf. Model 53, 1893–1904.

(80) Kadukova, M., and Grudinin, S. (2018) Docking of small molecules to farnesoid X receptors using AutoDock Vina with the Convex-PL potential: lessons learned from D3R Grand Challenge 2. J. Comput.-Aided Mol. Des. 32, 151–162.

(81) Kadukova, M., Chupin, V., and Grudinin, S. (2020) Docking rigid macrocycles using Convex-PL, AutoDock Vina, and RDKit in the D3R Grand Challenge 4. J. Comput.- Aided Mol. Des. 34, 191–200.

(82) Truchon, J.-F., and Bayly, C. I. (2007) Evaluating virtual screening methods: good and bad metrics for the “early recognition” problem. J. Chem. Inf. Model. 47, 488–508.

(83) Landrum, G. http://www.rdkit.org, RDKit: Open-source cheminformatics.

(84) Tran-Nguyen, V. K., Jacquemard, C., and Rognan, D. (2020) LIT-PCBA: An unbiased data set for machine learning and virtual screening. J. Chem. Inf. Model. 60, 4263–4273.

(85) Wang, S., Witek, J., Landrum, G. A., and Riniker, S. (2020) Improving Conformer Generation for Small Rings and Macrocycles Based on Distance Geometry and Experimental Torsional-Angle Preferences. J. Chem. Inf. Model 60, 2044–2058.

