## Supplementary Information for "Convex-PL^*R*^ – Revisiting affinity predictions and virtual screening using physics-informed machine learning"

#### Running header

Supplementary Information for Convex-PL$^R$ – Revisiting affinity predictions and virtual screening using physics-informed machine learning.

### Typization

Tables S1, S2 list the ligand and protein atom types used in Convex-PL$^{cof}$ and Convex-PL$^{R}$, and their chemical interpretation.

Table S1: 39 ligand atom types used in Convex-PL$^{R}$

| Type name | Description |
|---|---|
| C.ar6x | sp2 carbon in a 6-membered heteroaromatic ring |
| C.ar6 | sp2 carbon in a benzene ring |
| C.ar | sp2 carbon in other aromatics |
| C.sp1 | sp carbon |
| C.sp3 | sp3 carbon |
| C.co2 | sp2 carbon in COOH and COO- groups |
| C.guh | sp2 carbon in amidino and guanidino groups |
| C.sp2 | other sp2 carbon |
| N.arp | charged nitrogen in protonated aromatics |
| N.arA | other nitrogen in aromatic rings |
| N.1 | sp nitrogen |
| N.o2 | nitrogen in nitro groups |
| N.oh | nitrogen in hydroxamic acids and in hydroxylamines |
| N.amA | other sp2 nitrogen with three heavy atoms; nitrogen in sulfonamide, sulfonimide, carbon- or thionamide, aromatic amine, imide |
| N.guh | nitrogen in guanidino or amidino group |
| N.2p | other sp2 nitrogen with one heavy atom |
| N.2s | other sp2 nitrogen bonded to the two heavy atoms |
| N.3p | other sp3 nitrogen with one heavy atom bonded |
| N.3s | other sp3 nitrogen with 2 heavy atoms bonded |
| N.3t | other sp3 nitrogen with 3 heavy atoms bonded |
| N.4 | other sp3 nitrogen with 4 bonded atoms |
| O.3et | sp3 oxygen in ethers, esters, anhydrids, oxiran rings; sp3 oxygen with 2 heavy atoms bonded to at least one phosphor or sulfur |
| O.n | oxygen in nitro groups |
| O.3ac | sp3 oxygen in hydroxylamine or hydroxamic acid, COOH, CSOH, POOHOH, POOH or SOOOH |
| O.carb | sp2 oxygen in esters, anhydrids, carbonamides, acidhalogenides, COOH, POOHOH, POOH, SOOOH, and in other carbonyl groups |
| O.co2 | sp2 oxygen in OSO3-, SO3-, POO-, deprotonated sulfonamides, COO-, CSO-, OPO3H-, PO3H-, POO-, peroxo groups |
| O.3oh | hydroxyl group |
| O.ar | aromatic oxygen |
| S.r | aromatic sulfur; sulfur in a thiiran ring |

| | |
|---|---|
| S.o | sulfur in SO, SO3, CSO-, COS-, OSO3, OPO2SH-, PO2SH-, POS-, thionyl group, SO3-, OSO3- |
| S.o2 | sulfur in SO2 and sulfonamides |
| S.3 | other sulfur |
| P.o | phosphorus in groups with oxygen |
| P.3 | phosphorus in phosphiran rings; other sp3 phosphorus |
| F.0 | fluorine |
| Cl.0 | chlorine |
| Br.0 | bromine |
| I.0 | iodine |
| BSi | metalloids (B, Si, S, As) |

Table S2: 33 protein atom types used in Convex-PL$^R$

| Type name | Description |
|---|---|
| C.am | carbon of an amide group |
| C.3a | CA carbon |
| C.ar6 | carbon in a benzene ring |
| C.arp | carbon in protonated aromatics |
| C.arx | carbon in other aromatics |
| C.co2 | sp2 carbon in COOH and COO- groups |
| C.guh | sp2 carbon in amidino and guanidino groups |
| C.3p | sp3 terminal carbon |
| C.3s | sp3 aliphatic carbon with 2 heavy neighbours |
| C.3t | sp3 carbon with 3 heavy neighbours |
| C.3 | any other sp3 carbon |
| C.2 | any other sp2 carbon |
| N.amp | primary amide nitrogen (corresponds to a secondary amide in an actual peptide chain) |
| N.ams | primary amide nitrogen (corresponds to a tertiary amide in an actual peptide chain) |
| N.arp | protonated nitrogen in aromatics |
| N.ar2 | nitrogen in aromatics |
| N.guh | nitrogen in guanidino or amidino group |
| N.3p | terminal sp3 nitrogen |
| N.amA | any other flat nitrogen |
| N.3 | any other sp3 nitrogen |
| O.am | aromatic oxygen |
| O.carb | carboxyl oxygen |
| O.3oh | hydroxyl group |
| O.co2 | sp2 oxygen in OSO3-, SO3-, POO-, deprotonated sulfonamides |
| O.3ac | sp3 oxygen in hydroxylamine or hydroxamic acid, COOH, CSOH, POOHOH, POOH or SOOOH |

| | |
|---|---|
| O.3et | sp3 oxygen in ethers, esters, anhydrids, oxiran rings; sp3 oxygen with 2 heavy atoms bonded to at least one phosphor or sulfur |
| S.sh | terminal sulfur |
| S.o | sulfur in oxygen-containing groups |
| S.3 | other sulfur |
| Se | selene |
| P.o | phosphorus |
| Hal | halogen |
| Me | some metal atoms (Fe, Zn, Mg) – this type mostly represents heme Fe atom, as many hemes were available in the training set |

### Model training

#### Building the cofactor-containing dataset

Table S3: Number of PDB codes included and excluded from the linear regression training set.

| Condition | Number of complexes |
|---|---|
| PDBBind 2019 general set with avg RSCC and occupancy $< 0.8$, $\|\text{Z-score}\| > 15$ for any bond | 15,967 |
| Binding MOAD 2020 with avg RSCC and occupancy $< 0.8$, $\|\text{Z-score}\| > 15$ for any bond, no multiple ligands and badly formatted files | 3,445 |
| Complexes from PDB with modified residues, avg RSCC and occupancy $< 0.8$, receptor size $> 30$ residues, distance between the modified residue and ligand $< 5.2$Å, ligand is not a solvent/metal cluster. Each separate ligand forms a separate "complex" with the protein. | 1,783 |
| Complexes from PDB with cofactors, avg RSCC and occupancy $< 0.8$, distance between the cofactor and ligand $< 5.2$Å, ligand is not a solvent/metal cluster. Each ligand and cofactor are treated as a "ligand" and "protein" and vice versa, forming multiple training complexes. | 8,568 |
| Resulting number of complexes | 29,763 |

#### Selection criteria for the complexes from the PDBBind 2019 general set used in the linear regression model training

Table S4: Number of PDB codes included and excluded from the linear regression training set.

| Condition | Number of complexes satisfying the condition, excluding already filtered complexes |
|---|---|
| PDBBind 2019 general set | 17679 |
| CASF 2013 and 2016 | -373 |
| $\approx$ known binding constants | -116 |
| $>, <, \geq, \leq$ binding constants | -332 |
| "incorrect" label in the index file | -10 |
| $-\log K \geq 13$ | -14 |
| Convex-PL$^{cof}$ $\leq 0$ | -66 |

| | |
|---|---|
| other | -20 |
| Well correlated with Convex-PL$^{cof}$ | -4729 |
| Resulting number of complexes | 12019 |

#### Linear regression coefficients

Table S5: Feature weights. All features are scaled to the $[0, 1]$ interval.

| Feature | Weights Convex-PL$^{R}$ |
|---|---|
| Convex-PL$^{cof}$ | 7.28 |
| ligand flexibility | -2.20 |
| side-chain flexibility | -2.28 |

#### Interactions with the solvent

To account for the potentially underestimated interactions, we computed two types of descriptors. The first one were the solvent-accessible surface areas of atoms of different types. We have tested two ways of calculating these descriptors using the POWERSASA library[1,2] with a 1.4 Å probe atom radius. The first one was the computation of the usual ΔSASA buried upon binding. As an alternative, we computed SASAs only for those atoms that are considered to be interacting upon binding, i.e., are closer to each other than a cutoff distance that we set to 7Å. Both ways lead to similar results. We also tried different parametrizations of SASA-based descriptors, i.e. computed them for all protein and ligand atom types present in Convex-PL, grouped atoms by their chemical properties, or trained the model with only those groups of atom types that would penalize the number of contacts.

For those solvation interactions that were possibly not taken into account by the combination of the Convex-PL term and SASA, we decided to use a grid representation of solvent following the ideas from the SBROD protein quality assessment scoring function.[3] We first constructed three grids corresponding to the ligand, protein, and the whole complex centered on the ligand molecule with the size equal to the ligand size plus a 5 Å padding. For each molecule of the ligand, protein, and the complex, we removed grid nodes intersecting with the molecule within the 1.4 Å margins. We then collected statistics of distance distributions between the grid points and the atoms of the complex, protein, and ligand of different types. We wrote it to the *structure vectors* $x_{PL}^{sol}, x_{P}^{sol}$, and $x_{L}^{sol}$, correspondingly, and then computed the feature representing the difference of the solvation geometry between the bound and unbound states, $x^{sol} = x_{PL}^{sol} - (x_{P}^{sol} + x_{L}^{sol})$. Figure S1 shows an example of the solvent grid for 1gqs complex.

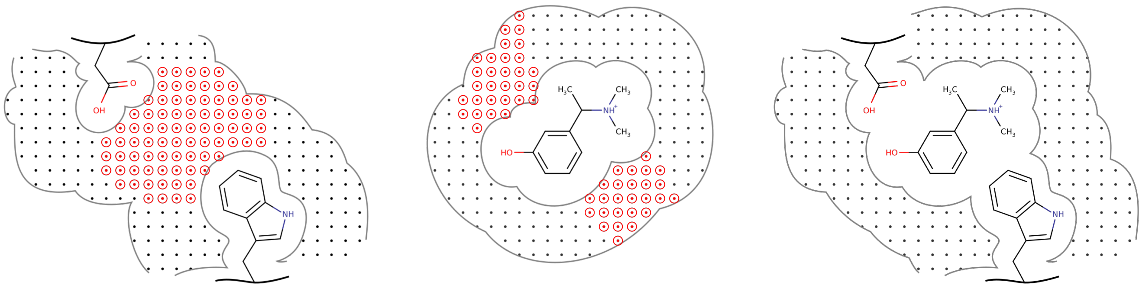

(a) (b) (c)

Figure S1: Schematic representation of solvent grids for a protein (a), ligand (b), and their complex (c). The difference between the complex grid and receptor and ligand grids is shown in red.

### CASF benchmarks

#### Evaluation of Convex-PL on the CASF benchmarks

Tables S6, S7, S8 list the results of Convex-PL, Convex-PL$^{cof}$, and Convex-PL$^{R}$ in the CASF-2013 and CASF-2016 benchmarks. Convex-PL pose prediction results in CASF-2013 are slightly better than those reported in[4], since we have updated the clash term.

Table S6: Success rates of finding native and near-native docking poses with RMSD smaller than 1 Å , 2 Å , and 3 Å among 1, 2, and 3 top-ranked structures scored with Convex-PL, Convex-PL$^{5.2A}$, and Convex-PL$^{R}$ in the CASF-2013 and CASF-2016 docking tests.

Convex-PL

| | CASF 2013 | | | | | | CASF-2016 | | | | | |
|---|---|---|---|---|---|---|---|---|---|---|---|---|
| | With natives | | | Without natives | | | With natives | | | Without natives | | |
| | Q1 | Q2 | Q3 | Q1 | Q2 | Q3 | Q1 | Q2 | Q3 | Q1 | Q2 | Q3 |
| Top 1 | 81.03 | 88.72 | 92.31 | 71.28 | 86.15 | 90.77 | 85.61 | 89.82 | 93.33 | 75.09 | 84.21 | 90.18 |
| Top 2 | 86.67 | 92.31 | 94.87 | 78.46 | 89.23 | 93.85 | 90.88 | 94.74 | 95.44 | 82.46 | 89.82 | 92.98 |
| Top 3 | 89.74 | 93.33 | 95.90 | 83.08 | 91.28 | 94.87 | 93.33 | 96.84 | 97.54 | 85.26 | 93.68 | 95.79 |

Convex-PL$^{R}$

| | CASF 2013 | | | | | | CASF-2016 | | | | | |
|---|---|---|---|---|---|---|---|---|---|---|---|---|
| | With natives | | | Without natives | | | With natives | | | Without natives | | |
| | Q1 | Q2 | Q3 | Q1 | Q2 | Q3 | Q1 | Q2 | Q3 | Q1 | Q2 | Q3 |
| Top 1 | 80.00 | 87.18 | 90.26 | 73.33 | 85.64 | 88.72 | 85.61 | 91.23 | 93.68 | 76.49 | 86.67 | 90.18 |
| Top 2 | 88.21 | 92.82 | 95.90 | 79.49 | 90.26 | 93.85 | 90.53 | 95.09 | 95.79 | 83.16 | 91.93 | 95.09 |
| Top 3 | 93.33 | 96.41 | 98.46 | 83.08 | 91.79 | 95.90 | 93.33 | 96.84 | 97.19 | 85.61 | 94.74 | 96.84 |

Convex-PL$^{cof}$

| | CASF 2013 | | | | | | CASF-2016 | | | | | |
|---|---|---|---|---|---|---|---|---|---|---|---|---|
| | With natives | | | Without natives | | | With natives | | | Without natives | | |
| | Q1 | Q2 | Q3 | Q1 | Q2 | Q3 | Q1 | Q2 | Q3 | Q1 | Q2 | Q3 |
| Top 1 | 81.54 | 87.69 | 90.77 | 75.38 | 85.13 | 88.21 | 85.96 | 91.58 | 92.98 | 77.54 | 86.67 | 89.82 |
| Top 2 | 87.18 | 92.82 | 95.38 | 78.97 | 89.23 | 93.33 | 90.88 | 95.79 | 96.14 | 83.86 | 91.93 | 94.74 |
| Top 3 | 92.82 | 95.90 | 97.44 | 84.10 | 91.28 | 95.38 | 93.68 | 96.84 | 97.54 | 86.67 | 94.39 | 96.49 |

Table S7: Scoring and ranking test results. $r_p$, $r_s$ correspond to the Pearson's and Spearman's correlation coefficients. In CASF-2013, the ranking performance is measured as a success rate of (high-level) correct ranking of all the three ligands binding the target protein, and (low-level) ranking the best complex as the top one.

| Scoring function | CASF-2013 | | | CASF-2016 | |
|---|---|---|---|---|---|
| | Scoring | Ranking | | Scoring | Ranking |
| | $r_p$ | "high" | "low" | $r_p$ | $r_s$ |
| Convex-PL | 0.574 | 60.0 | 70.8 | 0.607 | 0.584 |
| Convex-PL$^R$ | 0.618185 | 61.5 | 75.4 | 0.656994 | 0.610526 |
| Convex-PL$^{cof}$ | 0.609898 | 64.6 | 75.4 | 0.626039 | 0.615789 |

Table S8: CASF-2013 and CASF-2016 screening test results. Enrichment, best binder, and best target correspond to enrichment factors, a success rate of finding the strongest binding ligand for each protein target, and a success rate to find the strongest binding target for each ligand.

| Scoring function | CASF 2013 | | | | | | CASF-2016 | | | | | |
|---|---|---|---|---|---|---|---|---|---|---|---|---|
| | Enrichment | | | Best binder | | | Enrichment | | | Best binder | | |
| | Top | | | Top | | | Top | | | Top | | |
| | 1% | 5% | 10% | 1% | 5% | 10% | 1% | 5% | 10% | 1% | 5% | 10% |
| Convex-PL | 9.5 | 4.7 | 3.1 | 27.7 | 55.4 | 64.6 | 4.6 | 3.1 | 2.7 | 14.0 | 35.1 | 61.4 |
| Convex-PL$^R$ | 26.2 | 7.0 | 4.3 | 73.8 | 80.0 | 84.6 | 12.5 | 5.1 | 3.5 | 35.1 | 54.4 | 70.2 |
| Convex-PL$^{cof}$ | 18.8 | 6.4 | 4.0 | 53.8 | 73.8 | 80.0 | 8.0 | 4.2 | 3.3 | 28.1 | 45.6 | 66.7 |

### D3R benchmark

Tables S10, S11, S12, S13, S14, S9 list the results of Convex-PL, Convex-PL$^R$, Convex-PL$^{cof}$, KORP-PL$^w$, AutoDock Vina, and ΔSAS in the benchmark obtained from the D3R Challenges structures, user submissions, and binding constants.

Table S9: Correlation between the predicted binding affinity of the native poses from the D3R Grand Challenge 2, Grand Challenge 3, and Grand Challenge 4, and the experimentally obtained binding constants. $r_p$, $r_s$ correspond to the Pearson's and Spearman's correlation coefficients, $\tau$ denotes the Kendall's $\tau$ coefficient.

Scoring test, native structures

| Scoring function | Correlation coefficients | | | | | | | | |
|---|---|---|---|---|---|---|---|---|---|
| | GC2 | | | GC3 | | | GC4 | | |
| | $r_p$ | $r_s$ | $\tau$ | $r_p$ | $r_s$ | $\tau$ | $r_p$ | $r_s$ | $\tau$ |
| Convex-PL | 0.238 | 0.159 | 0.123 | 0.148 | 0.092 | 0.065 | 0.165 | 0.071 | 0.067 |
| Convex-PL$^R$ | 0.489 | 0.414 | 0.281 | 0.169 | 0.134 | 0.124 | 0.244 | 0.011 | 0.033 |
| Convex-PL$^{cof}$ | 0.531 | 0.455 | 0.294 | 0.417 | 0.428 | 0.349 | 0.263 | 0.008 | 0.033 |
| KORP-PL$^w$ | 0.564 | 0.587 | 0.415 | 0.014 | -0.008 | -0.006 | 0.343 | 0.376 | 0.233 |
| AutoDock Vina | 0.249 | 0.187 | 0.129 | 0.195 | 0.112 | 0.075 | 0.407 | 0.515 | 0.333 |
| ΔSAS | 0.49 | 0.37 | 0.24 | 0.44 | 0.39 | 0.33 | 0.53 | 0.17 | 0.12 |

Table S10: Success rates of finding native and near-native docking poses with RMSD smaller than 1 Å , 2 Å , and 3 Å in the 1%, 5%, and 10% of the structures taken from the D3R Grand Challenge 2 user submissions.

Pose prediction test, D3R Grand Challenge 2

| Scoring function | With natives | | | | | | | | | Without natives | | | | | | | | |
|---|---|---|---|---|---|---|---|---|---|---|---|---|---|---|---|---|---|---|
| | Q1 | | | Q2 | | | Q3 | | | Q1 | | | Q2 | | | Q3 | | |
| | top, % | | | top, % | | | top, % | | | top, % | | | top, % | | | top, % | | |
| | 1 | 5 | 10 | 1 | 5 | 10 | 1 | 5 | 10 | 1 | 5 | 10 | 1 | 5 | 10 | 1 | 5 | 10 |
| Convex-PL | 0.51 | 0.8 | 0.94 | 0.71 | 0.91 | 0.97 | 0.83 | 0.94 | 0.97 | 0.62 | 0.83 | 0.83 | 0.7 | 0.85 | 0.88 | 0.79 | 0.88 | 0.88 |
| Convex-PL$^R$ | 0.54 | 0.86 | 0.94 | 0.74 | 0.91 | 0.97 | 0.83 | 0.97 | 1.0 | 0.71 | 0.83 | 0.92 | 0.76 | 0.85 | 0.88 | 0.82 | 0.88 | 0.88 |
| Convex-PL$^{cof}$ | 0.54 | 0.86 | 0.97 | 0.74 | 0.91 | 1.0 | 0.83 | 0.97 | 1.0 | 0.71 | 0.83 | 0.92 | 0.76 | 0.85 | 0.85 | 0.82 | 0.88 | 0.88 |
| KORP-PL$^w$ | 0.49 | 0.66 | 0.71 | 0.6 | 0.66 | 0.77 | 0.66 | 0.77 | 0.86 | 0.67 | 0.92 | 0.92 | 0.61 | 0.67 | 0.76 | 0.65 | 0.79 | 0.88 |
| AutoDock Vina | 0.51 | 0.69 | 0.8 | 0.66 | 0.86 | 0.89 | 0.71 | 0.91 | 0.94 | 0.71 | 0.88 | 0.88 | 0.67 | 0.88 | 0.88 | 0.71 | 0.91 | 0.91 |
| ΔSAS | 0.29 | 0.57 | 0.71 | 0.6 | 0.8 | 0.86 | 0.63 | 0.86 | 0.91 | 0.42 | 0.75 | 0.88 | 0.64 | 0.82 | 0.85 | 0.65 | 0.88 | 0.91 |

Table S11: Success rates of finding native and near-native docking poses with RMSD smaller than 1 Å , 2 Å , and 3 Å in the 1%, 5%, and 10% of the structures taken from the D3R Grand Challenge 3 Stage 1a user submissions.

Pose prediction test, D3R Grand Challenge 3

| Scoring function | With natives | | | | | | | | | Without natives | | | | | | | | |
|---|---|---|---|---|---|---|---|---|---|---|---|---|---|---|---|---|---|---|
| | Q1 | | | Q2 | | | Q3 | | | Q1 | | | Q2 | | | Q3 | | |
| | top, % | | | top, % | | | top, % | | | top, % | | | top, % | | | top, % | | |
| | 1 | 5 | 10 | 1 | 5 | 10 | 1 | 5 | 10 | 1 | 5 | 10 | 1 | 5 | 10 | 1 | 5 | 10 |
| Convex-PL | 0.04 | 0.17 | 0.54 | 0.08 | 0.42 | 0.79 | 0.25 | 0.71 | 0.92 | 0.08 | 0.17 | 0.5 | 0.09 | 0.39 | 0.74 | 0.25 | 0.71 | 0.88 |
| Convex-PL$^R$ | 0.04 | 0.46 | 0.58 | 0.17 | 0.67 | 0.88 | 0.46 | 0.83 | 1.0 | 0.0 | 0.25 | 0.5 | 0.13 | 0.52 | 0.74 | 0.42 | 0.75 | 0.92 |
| Convex-PL$^{cof}$ | 0.08 | 0.5 | 0.58 | 0.17 | 0.71 | 0.88 | 0.29 | 0.88 | 0.96 | 0.0 | 0.08 | 0.5 | 0.09 | 0.57 | 0.78 | 0.25 | 0.79 | 0.88 |
| KORP-PL$^w$ | 0.29 | 0.54 | 0.79 | 0.5 | 0.83 | 0.92 | 0.62 | 0.96 | 1 | 0.42 | 0.58 | 0.75 | 0.48 | 0.78 | 0.87 | 0.58 | 0.88 | 0.96 |
| AutoDock Vina | 0 | 0 | 0 | 0.04 | 0.04 | 0.08 | 0.04 | 0.12 | 0.12 | 0 | 0 | 0 | 0.04 | 0.04 | 0.09 | 0.04 | 0.12 | 0.12 |
| ΔSAS | 0 | 0 | 0 | 0 | 0.04 | 0.04 | 0.08 | 0.12 | 0.25 | 0 | 0 | 0 | 0 | 0.04 | 0.04 | 0.08 | 0.12 | 0.25 |

Table S12: Success rates of finding native and near-native docking poses with RMSD smaller than 1 Å , 2 Å , and 3 Å in the 1%, 5%, and 10% of the structures taken from the D3R Grand Challenge 3 Stage 1b user submissions.

Pose prediction test, D3R Grand Challenge 3

| Scoring function | With natives | | | | | | | | | Without natives | | | | | | | | |
|---|---|---|---|---|---|---|---|---|---|---|---|---|---|---|---|---|---|---|
| | Q1 | | | Q2 | | | Q3 | | | Q1 | | | Q2 | | | Q3 | | |
| | top, % | | | top, % | | | top, % | | | top, % | | | top, % | | | top, % | | |
| | 1 | 5 | 10 | 1 | 5 | 10 | 1 | 5 | 10 | 1 | 5 | 10 | 1 | 5 | 10 | 1 | 5 | 10 |
| Convex-PL | 0.0 | 0.08 | 0.62 | 0.0 | 0.17 | 0.67 | 0.04 | 0.29 | 0.75 | 0.0 | 0.07 | 0.86 | 0.0 | 0.12 | 0.67 | 0.04 | 0.25 | 0.75 |
| Convex-PL$^R$ | 0.12 | 0.71 | 0.71 | 0.21 | 0.75 | 0.83 | 0.25 | 0.83 | 0.88 | 0.0 | 0.57 | 0.86 | 0.08 | 0.5 | 0.71 | 0.12 | 0.67 | 0.83 |
| Convex-PL$^{cof}$ | 0.0 | 0.67 | 0.79 | 0.12 | 0.79 | 0.92 | 0.17 | 0.83 | 0.96 | 0.0 | 0.57 | 0.93 | 0.12 | 0.54 | 0.75 | 0.17 | 0.62 | 0.83 |
| KORP-PL$^w$ | 0.42 | 0.75 | 0.88 | 0.54 | 1 | 1 | 0.67 | 1 | 1 | 0.5 | 0.71 | 0.79 | 0.46 | 0.96 | 0.96 | 0.62 | 0.96 | 1 |
| AutoDock Vina | 0 | 0.08 | 0.12 | 0.04 | 0.12 | 0.21 | 0.04 | 0.12 | 0.33 | 0 | 0.07 | 0.07 | 0.04 | 0.12 | 0.17 | 0.04 | 0.12 | 0.29 |
| ΔSAS | 0 | 0 | 0.12 | 0 | 0 | 0.29 | 0 | 0.12 | 0.33 | 0 | 0 | 0.14 | 0 | 0 | 0.25 | 0 | 0.12 | 0.33 |

Table S13: Success rates of finding native and near-native docking poses with RMSD smaller than 1 Å , 2 Å , and 3 Å in the 1%, 5%, and 10% of the structures taken from the D3R Grand Challenge 4 Stage 1a user submissions.

Pose prediction test, D3R Grand Challenge 4

| Scoring function | With natives | | | | | | | | | Without natives | | | | | | | | |
|---|---|---|---|---|---|---|---|---|---|---|---|---|---|---|---|---|---|---|
| | Q1 | | | Q2 | | | Q3 | | | Q1 | | | Q2 | | | Q3 | | |
| | top, % | | | top, % | | | top, % | | | top, % | | | top, % | | | top, % | | |
| | 1 | 5 | 10 | 1 | 5 | 10 | 1 | 5 | 10 | 1 | 5 | 10 | 1 | 5 | 10 | 1 | 5 | 10 |
| Convex-PL | 0.8 | 1.0 | 1.0 | 1 | 1 | 1 | 1 | 1 | 1 | 0.7 | 0.95 | 0.95 | 0.9 | 1.0 | 1.0 | 0.95 | 1.0 | 1.0 |
| Convex-PL$^R$ | 0.55 | 1.0 | 1.0 | 0.85 | 1.0 | 1.0 | 0.85 | 1.0 | 1.0 | 0.35 | 0.9 | 0.95 | 0.85 | 1.0 | 1.0 | 0.85 | 1.0 | 1.0 |
| Convex-PL$^{cof}$ | 0.75 | 1.0 | 1.0 | 0.95 | 1.0 | 1.0 | 0.95 | 1.0 | 1.0 | 0.45 | 0.95 | 0.95 | 0.9 | 1.0 | 1.0 | 0.9 | 1.0 | 1.0 |
| KORP-PL$^w$ | 0.95 | 1 | 1 | 1 | 1 | 1 | 1 | 1 | 1 | 0.65 | 0.9 | 1 | 1 | 1 | 1 | 1 | 1 | 1 |
| AutoDock Vina | 0.5 | 0.95 | 1 | 0.95 | 1 | 1 | 0.95 | 1 | 1 | 0.5 | 0.95 | 1 | 0.95 | 1 | 1 | 0.95 | 1 | 1 |
| ΔSAS | 0.4 | 0.75 | 0.9 | 0.8 | 1 | 1 | 0.9 | 1 | 1 | 0.4 | 0.75 | 0.9 | 0.8 | 1 | 1 | 0.9 | 1 | 1 |

Table S14: Success rates of finding native and near-native docking poses with RMSD smaller than 1 Å , 2 Å , and 3 Å in the 1%, 5%, and 10% of the structures taken from the D3R Grand Challenge 4 Stage 1b user submissions.

Pose prediction test, D3R Grand Challenge 4

| Scoring function | With natives | | | | | | | | | Without natives | | | | | | | | |
|---|---|---|---|---|---|---|---|---|---|---|---|---|---|---|---|---|---|---|
| | Q1 | | | Q2 | | | Q3 | | | Q1 | | | Q2 | | | Q3 | | |
| | top, % | | | top, % | | | top, % | | | top, % | | | top, % | | | top, % | | |
| | 1 | 5 | 10 | 1 | 5 | 10 | 1 | 5 | 10 | 1 | 5 | 10 | 1 | 5 | 10 | 1 | 5 | 10 |
| Convex-PL | 0.65 | 0.95 | 1.0 | 1 | 1 | 1 | 1 | 1 | 1 | 0.65 | 0.95 | 0.95 | 1 | 1 | 1 | 1 | 1 | 1 |
| Convex-PL$^R$ | 0.85 | 0.95 | 1.0 | 1.0 | 1.0 | 1.0 | 1.0 | 1.0 | 1.0 | 0.7 | 0.9 | 0.95 | 1.0 | 1.0 | 1.0 | 1.0 | 1.0 | 1.0 |
| Convex-PL$^{cof}$ | 0.85 | 0.95 | 1.0 | 1.0 | 1.0 | 1.0 | 1.0 | 1.0 | 1.0 | 0.75 | 0.9 | 0.95 | 1.0 | 1.0 | 1.0 | 1.0 | 1.0 | 1.0 |
| KORP-PL$^w$ | 1 | 1 | 1 | 1 | 1 | 1 | 1 | 1 | 1 | 0.65 | 1 | 1 | 0.95 | 1 | 1 | 1 | 1 | 1 |
| AutoDock Vina | 0.55 | 1 | 1 | 0.95 | 1 | 1 | 1 | 1 | 1 | 0.55 | 1 | 1 | 0.95 | 1 | 1 | 1 | 1 | 1 |
| ΔSAS | 0.15 | 0.7 | 0.8 | 0.8 | 1 | 1 | 0.8 | 1 | 1 | 0.15 | 0.7 | 0.8 | 0.8 | 1 | 1 | 0.8 | 1 | 1 |

### DUD benchmark

Table S15: A list of 9 DUD targets with co-factors.

| DUD targets with co-factors |
|---|
| alr2, comt, gart, gpb, pnp, sahh, tk, dhfr, inha |

Table S16: ROC AUC, BEDROC, and 5% EF computed for AutoDock Vina, Vinardo, Convex-PL, Convex-PL$^R$, Convex-PL$^{cof}$, and KORP-PL$^w$ in the DUD benchmark.

| | ROC AUC | | | | | | EF5% | | | | | | BEDROC$_{\alpha=20}$ | | | | | |
|---|---|---|---|---|---|---|---|---|---|---|---|---|---|---|---|---|---|---|
| Target | AutoDock Vina | Vinardo | Convex-PL | Convex-PL$^R$ | Convex-PL$^{cof}$ | KORP-PL$^w$ | AutoDock Vina | Vinardo | Convex-PL | Convex-PL$^R$ | Convex-PL$^{cof}$ | KORP-PL$^w$ | AutoDock Vina | Vinardo | Convex-PL | Convex-PL$^R$ | Convex-PL$^{cof}$ | KORP-PL$^w$ |
| ace | 0.373 | 0.432 | 0.451 | 0.593 | 0.574 | 0.741 | 1.633 | 2.041 | 4.082 | 6.531 | 6.939 | 7.347 | 0.116 | 0.136 | 0.202 | 0.379 | 0.397 | 0.437 |
| ache | 0.663 | 0.678 | 0.568 | 0.504 | 0.531 | 0.418 | 3.738 | 4.299 | 2.617 | 2.430 | 2.804 | 0.748 | 0.236 | 0.259 | 0.143 | 0.119 | 0.157 | 0.053 |
| ada | 0.503 | 0.599 | 0.224 | 0.665 | 0.628 | 0.670 | 0.000 | 2.564 | 0.000 | 2.051 | 1.538 | 4.103 | 0.013 | 0.128 | 0.004 | 0.140 | 0.120 | 0.220 |
| ampc | 0.313 | 0.441 | 0.461 | 0.668 | 0.673 | 0.907 | 0.952 | 0.952 | 0.000 | 0.000 | 0.000 | 16.191 | 0.029 | 0.059 | 0.016 | 0.038 | 0.037 | 0.824 |
| ar | 0.792 | 0.735 | 0.655 | 0.647 | 0.614 | 0.850 | 10.886 | 7.595 | 5.316 | 5.316 | 4.304 | 5.823 | 0.560 | 0.445 | 0.275 | 0.266 | 0.239 | 0.328 |
| cdk2 | 0.542 | 0.590 | 0.494 | 0.610 | 0.558 | 0.826 | 2.500 | 4.722 | 4.167 | 3.333 | 3.333 | 5.833 | 0.176 | 0.261 | 0.237 | 0.193 | 0.201 | 0.348 |
| cox1 | 0.675 | 0.673 | 0.456 | 0.583 | 0.574 | 0.571 | 7.200 | 8.000 | 0.800 | 0.800 | 0.800 | 2.400 | 0.389 | 0.362 | 0.072 | 0.145 | 0.117 | 0.160 |
| cox2 | 0.905 | 0.894 | 0.542 | 0.865 | 0.825 | 0.526 | 13.239 | 12.864 | 0.376 | 9.859 | 8.263 | 1.080 | 0.700 | 0.685 | 0.037 | 0.578 | 0.486 | 0.063 |
| egfr | 0.582 | 0.719 | 0.785 | 0.828 | 0.774 | 0.949 | 2.484 | 3.832 | 7.116 | 5.979 | 6.526 | 14.400 | 0.132 | 0.228 | 0.396 | 0.350 | 0.351 | 0.743 |
| er_agonist | 0.794 | 0.806 | 0.597 | 0.770 | 0.707 | 0.891 | 8.955 | 9.254 | 0.895 | 8.060 | 6.269 | 8.955 | 0.459 | 0.493 | 0.082 | 0.421 | 0.333 | 0.456 |
| er_antagonist | 0.686 | 0.697 | 0.720 | 0.783 | 0.731 | 0.832 | 4.615 | 7.179 | 5.128 | 8.205 | 7.179 | 10.769 | 0.276 | 0.352 | 0.271 | 0.435 | 0.396 | 0.551 |
| fgfr1 | 0.358 | 0.393 | 0.532 | 0.585 | 0.546 | 0.885 | 0.500 | 0.833 | 1.667 | 1.833 | 2.833 | 12.833 | 0.030 | 0.057 | 0.080 | 0.113 | 0.145 | 0.671 |
| fxa | 0.662 | 0.585 | 0.687 | 0.805 | 0.750 | 0.953 | 1.781 | 2.466 | 2.329 | 6.849 | 6.027 | 14.931 | 0.118 | 0.149 | 0.151 | 0.360 | 0.331 | 0.704 |
| gr | 0.539 | 0.473 | 0.516 | 0.455 | 0.434 | 0.736 | 2.308 | 1.795 | 1.795 | 1.282 | 1.282 | 4.359 | 0.141 | 0.111 | 0.109 | 0.093 | 0.077 | 0.279 |
| hivpr | 0.729 | 0.830 | 0.559 | 0.762 | 0.701 | 0.872 | 3.871 | 8.065 | 3.548 | 7.097 | 7.742 | 6.452 | 0.260 | 0.397 | 0.165 | 0.376 | 0.406 | 0.425 |
| hivrt | 0.643 | 0.634 | 0.559 | 0.617 | 0.619 | 0.715 | 4.651 | 4.186 | 2.326 | 4.186 | 4.651 | 6.046 | 0.257 | 0.223 | 0.141 | 0.222 | 0.247 | 0.335 |
| hmga | 0.660 | 0.803 | 0.463 | 0.550 | 0.573 | 0.776 | 3.429 | 4.000 | 0.571 | 6.857 | 8.571 | 8.000 | 0.181 | 0.229 | 0.036 | 0.365 | 0.467 | 0.427 |
| hsp90 | 0.568 | 0.776 | 0.269 | 0.673 | 0.633 | 0.825 | 0.000 | 4.324 | 0.000 | 0.000 | 1.622 | 4.324 | 0.040 | 0.200 | 0.019 | 0.042 | 0.127 | 0.275 |
| mr | 0.811 | 0.807 | 0.772 | 0.815 | 0.806 | 0.902 | 14.667 | 14.667 | 5.333 | 12.000 | 9.333 | 10.667 | 0.674 | 0.647 | 0.253 | 0.592 | 0.531 | 0.515 |
| na | 0.375 | 0.373 | 0.493 | 0.832 | 0.828 | 0.834 | 0.000 | 0.000 | 0.000 | 3.674 | 3.674 | 13.061 | 0.003 | 0.009 | 0.007 | 0.240 | 0.235 | 0.635 |
| p38 | 0.613 | 0.627 | 0.601 | 0.581 | 0.565 | 0.450 | 2.335 | 1.366 | 1.630 | 1.674 | 1.718 | 1.894 | 0.179 | 0.127 | 0.117 | 0.124 | 0.130 | 0.123 |
| parp | 0.711 | 0.603 | 0.691 | 0.772 | 0.744 | 0.878 | 4.000 | 2.286 | 2.286 | 8.571 | 8.000 | 9.143 | 0.218 | 0.113 | 0.120 | 0.416 | 0.399 | 0.516 |
| pde5 | 0.706 | 0.757 | 0.489 | 0.732 | 0.699 | 0.806 | 5.000 | 5.000 | 2.727 | 6.818 | 6.136 | 8.182 | 0.322 | 0.359 | 0.188 | 0.429 | 0.412 | 0.498 |
| pdgfrb | 0.326 | 0.372 | 0.360 | 0.347 | 0.344 | 0.549 | 1.294 | 1.294 | 1.059 | 1.294 | 1.294 | 5.882 | 0.082 | 0.079 | 0.076 | 0.078 | 0.078 | 0.332 |
| ppar | 0.743 | 0.925 | 0.925 | 0.941 | 0.936 | 0.920 | 4.471 | 14.823 | 16.000 | 17.412 | 17.177 | 14.353 | 0.222 | 0.740 | 0.798 | 0.878 | 0.875 | 0.721 |
| pr | 0.444 | 0.397 | 0.297 | 0.486 | 0.439 | 0.648 | 1.482 | 0.000 | 0.741 | 0.000 | 0.000 | 2.222 | 0.063 | 0.024 | 0.039 | 0.015 | 0.007 | 0.178 |
| rxr | 0.926 | 0.916 | 0.847 | 0.850 | 0.842 | 0.948 | 16.000 | 14.000 | 6.000 | 12.000 | 10.000 | 15.000 | 0.786 | 0.745 | 0.288 | 0.623 | 0.551 | 0.659 |
| src | 0.642 | 0.712 | 0.612 | 0.773 | 0.713 | 0.880 | 1.761 | 4.780 | 3.774 | 6.667 | 5.660 | 14.465 | 0.119 | 0.251 | 0.194 | 0.360 | 0.327 | 0.762 |
| thrombin | 0.691 | 0.516 | 0.707 | 0.724 | 0.716 | 0.815 | 4.722 | 2.500 | 5.000 | 7.778 | 9.444 | 12.222 | 0.281 | 0.134 | 0.307 | 0.357 | 0.451 | 0.577 |
| trypsin | 0.727 | 0.514 | 0.790 | 0.917 | 0.877 | 0.941 | 2.041 | 4.082 | 9.388 | 10.612 | 9.796 | 12.245 | 0.157 | 0.177 | 0.449 | 0.544 | 0.475 | 0.539 |
| vegfr2 | 0.707 | 0.721 | 0.606 | 0.744 | 0.695 | 0.885 | 7.500 | 5.909 | 6.591 | 3.864 | 4.318 | 10.227 | 0.407 | 0.352 | 0.326 | 0.241 | 0.282 | 0.558 |
| alr2 | 0.707 | 0.650 | 0.386 | 0.554 | 0.484 | 0.454 | 6.154 | 4.615 | 1.538 | 3.846 | 4.615 | 0.769 | 0.268 | 0.263 | 0.069 | 0.166 | 0.276 | 0.045 |
| comt | 0.553 | 0.524 | 0.362 | 0.580 | 0.413 | 0.380 | 5.455 | 5.455 | 5.455 | 5.455 | 5.455 | 0.000 | 0.287 | 0.357 | 0.294 | 0.318 | 0.320 | 0.065 |
| dhfr | 0.812 | 0.875 | 0.706 | 0.661 | 0.553 | 0.616 | 4.585 | 6.829 | 2.780 | 4.488 | 2.634 | 3.366 | 0.324 | 0.458 | 0.207 | 0.294 | 0.180 | 0.231 |
| gart | 0.733 | 0.860 | 0.913 | 0.939 | 0.928 | 0.913 | 0.000 | 3.000 | 4.500 | 10.000 | 8.500 | 8.500 | 0.056 | 0.238 | 0.341 | 0.605 | 0.577 | 0.492 |
| gpb | 0.682 | 0.659 | 0.347 | 0.592 | 0.562 | 0.920 | 3.846 | 1.923 | 0.000 | 0.385 | 0.769 | 13.846 | 0.174 | 0.101 | 0.005 | 0.054 | 0.047 | 0.672 |
| inha | 0.561 | 0.518 | 0.460 | 0.534 | 0.529 | 0.561 | 6.046 | 1.861 | 1.628 | 3.023 | 4.884 | 3.023 | 0.324 | 0.127 | 0.108 | 0.170 | 0.281 | 0.160 |
| pnp | 0.623 | 0.644 | 0.322 | 0.637 | 0.471 | 0.995 | 0.000 | 4.000 | 0.000 | 0.800 | 0.400 | 16.800 | 0.081 | 0.194 | 0.006 | 0.043 | 0.048 | 0.943 |
| sahh | 0.829 | 0.769 | 0.559 | 0.694 | 0.701 | 0.558 | 10.303 | 7.879 | 0.000 | 3.636 | 3.636 | 0.000 | 0.528 | 0.374 | 0.013 | 0.185 | 0.152 | 0.012 |
| tk | 0.527 | 0.541 | 0.544 | 0.499 | 0.483 | 0.772 | 0.000 | 0.909 | 0.909 | 0.000 | 0.000 | 5.455 | 0.018 | 0.063 | 0.067 | 0.022 | 0.034 | 0.334 |
| mean | 0.636 | 0.651 | 0.558 | 0.679 | 0.644 | 0.764 | 4.360 | 4.904 | 3.002 | 5.117 | 4.953 | 7.898 | 0.242 | 0.268 | 0.168 | 0.285 | 0.283 | 0.422 |
| median | 0.663 | 0.655 | 0.552 | 0.667 | 0.631 | 0.825 | 3.792 | 4.134 | 2.306 | 4.337 | 4.633 | 7.673 | 0.200 | 0.228 | 0.131 | 0.253 | 0.278 | 0.432 |

Table S17: 1%, and 0.5% EF computed for AutoDock Vina, Vinardo, Convex-PL, Convex-PL$^R$, Convex-PL$^{cof}$, and KORP-PL$^w$ in the DUD benchmark.

| | EF1% | | | | | | EF0.5% | | | | | |
|---|---|---|---|---|---|---|---|---|---|---|---|---|
| Target | AutoDock Vina | Vinardo | Convex-PL | Convex-PL$^R$ | Convex-PL$^{cof}$ | KORP-PL$^w$ | AutoDock Vina | Vinardo | Convex-PL | Convex-PL$^R$ | Convex-PL$^{cof}$ | KORP-PL$^w$ |
| ace | 4.082 | 8.163 | 10.204 | 16.326 | 20.408 | 24.490 | 4.082 | 12.245 | 12.245 | 28.571 | 36.735 | 36.735 |
| ache | 1.869 | 4.673 | 4.673 | 4.673 | 8.411 | 0.935 | 0.000 | 1.869 | 3.738 | 9.346 | 9.346 | 0.000 |
| ada | 0.000 | 2.564 | 0.000 | 2.564 | 2.564 | 0.000 | 0.000 | 5.128 | 0.000 | 0.000 | 5.128 | 0.000 |
| ampc | 0.000 | 4.762 | 0.000 | 0.000 | 0.000 | 33.333 | 0.000 | 0.000 | 0.000 | 0.000 | 0.000 | 38.095 |
| ar | 17.721 | 21.519 | 11.392 | 7.595 | 7.595 | 2.532 | 22.785 | 27.848 | 17.721 | 7.595 | 7.595 | 2.532 |
| cdk2 | 5.556 | 11.111 | 9.722 | 2.778 | 6.944 | 9.722 | 8.333 | 11.111 | 16.667 | 5.556 | 8.333 | 13.889 |
| cox1 | 12.000 | 8.000 | 0.000 | 4.000 | 4.000 | 4.000 | 16.000 | 8.000 | 0.000 | 8.000 | 0.000 | 8.000 |
| cox2 | 25.117 | 26.056 | 0.000 | 24.883 | 21.596 | 0.939 | 25.822 | 30.047 | 0.000 | 26.291 | 23.005 | 0.469 |
| egfr | 2.947 | 7.158 | 17.053 | 10.737 | 11.579 | 26.737 | 3.368 | 7.158 | 23.579 | 14.737 | 16.421 | 28.632 |
| er_agonist | 16.418 | 16.418 | 0.000 | 19.403 | 16.418 | 11.940 | 11.940 | 20.895 | 0.000 | 20.895 | 20.895 | 20.895 |
| er_antagonist | 10.256 | 12.820 | 7.692 | 15.385 | 15.385 | 23.077 | 10.256 | 20.513 | 0.000 | 10.256 | 10.256 | 30.769 |
| fgfr1 | 0.833 | 1.667 | 0.000 | 1.667 | 0.833 | 30.000 | 0.000 | 1.667 | 0.000 | 0.000 | 0.000 | 38.333 |
| fxa | 0.685 | 6.164 | 4.795 | 11.644 | 12.329 | 17.808 | 1.370 | 5.479 | 5.479 | 15.069 | 17.808 | 17.808 |
| gr | 7.692 | 5.128 | 2.564 | 5.128 | 3.846 | 10.256 | 10.256 | 7.692 | 0.000 | 5.128 | 2.564 | 15.385 |
| hivpr | 6.452 | 11.290 | 3.226 | 11.290 | 11.290 | 16.129 | 6.452 | 6.452 | 6.452 | 16.129 | 19.355 | 12.903 |
| hivrt | 11.628 | 6.977 | 6.977 | 9.302 | 9.302 | 13.954 | 13.954 | 4.651 | 9.302 | 13.954 | 13.954 | 18.605 |
| hmga | 2.857 | 2.857 | 0.000 | 17.143 | 37.143 | 28.571 | 0.000 | 0.000 | 0.000 | 28.571 | 45.714 | 28.571 |
| hsp90 | 0.000 | 0.000 | 0.000 | 0.000 | 2.703 | 8.108 | 0.000 | 0.000 | 0.000 | 0.000 | 5.405 | 16.216 |
| mr | 26.667 | 26.667 | 0.000 | 26.667 | 20.000 | 6.667 | 26.667 | 13.333 | 0.000 | 13.333 | 26.667 | 13.333 |
| na | 0.000 | 0.000 | 0.000 | 12.245 | 12.245 | 16.326 | 0.000 | 0.000 | 0.000 | 16.326 | 12.245 | 28.571 |
| p38 | 1.762 | 2.423 | 1.982 | 2.203 | 2.203 | 3.744 | 0.441 | 4.405 | 3.084 | 3.084 | 3.965 | 3.524 |
| parp | 2.857 | 0.000 | 0.000 | 17.143 | 20.000 | 20.000 | 0.000 | 0.000 | 0.000 | 22.857 | 22.857 | 11.429 |
| pde5 | 11.364 | 13.636 | 7.955 | 15.909 | 15.909 | 22.727 | 18.182 | 22.727 | 9.091 | 22.727 | 22.727 | 22.727 |
| pdgfrb | 6.471 | 5.882 | 4.118 | 5.294 | 5.294 | 17.059 | 11.765 | 11.765 | 8.235 | 8.235 | 8.235 | 24.706 |
| ppar | 3.529 | 27.059 | 28.235 | 32.941 | 32.941 | 27.059 | 2.353 | 28.235 | 30.588 | 35.294 | 35.294 | 30.588 |
| pr | 0.000 | 0.000 | 3.704 | 0.000 | 0.000 | 11.111 | 0.000 | 0.000 | 0.000 | 0.000 | 0.000 | 14.815 |
| rxr | 40.000 | 35.000 | 0.000 | 35.000 | 20.000 | 20.000 | 40.000 | 40.000 | 0.000 | 40.000 | 30.000 | 20.000 |
| src | 2.516 | 5.660 | 4.402 | 12.579 | 15.723 | 38.994 | 0.000 | 7.547 | 3.774 | 15.094 | 25.157 | 38.994 |
| thrombin | 11.111 | 2.778 | 0.000 | 2.778 | 8.333 | 8.333 | 19.444 | 0.000 | 0.000 | 2.778 | 5.556 | 5.556 |
| trypsin | 2.041 | 6.122 | 0.000 | 12.245 | 8.163 | 0.000 | 4.082 | 8.163 | 0.000 | 12.245 | 8.163 | 0.000 |
| vegfr2 | 20.454 | 14.773 | 14.773 | 6.818 | 11.364 | 23.864 | 27.273 | 15.909 | 11.364 | 4.545 | 15.909 | 27.273 |
| alr2 | 3.846 | 11.539 | 0.000 | 0.000 | 15.385 | 0.000 | 7.692 | 23.077 | 0.000 | 0.000 | 15.385 | 0.000 |
| comt | 9.091 | 27.273 | 18.182 | 27.273 | 27.273 | 0.000 | 18.182 | 36.364 | 36.364 | 36.364 | 36.364 | 0.000 |
| dhfr | 9.512 | 9.756 | 4.634 | 9.268 | 6.829 | 6.829 | 9.756 | 9.756 | 7.805 | 15.122 | 11.220 | 8.293 |
| gart | 0.000 | 0.000 | 2.500 | 12.500 | 15.000 | 5.000 | 0.000 | 0.000 | 5.000 | 20.000 | 15.000 | 5.000 |
| gpb | 1.923 | 0.000 | 0.000 | 0.000 | 0.000 | 25.000 | 3.846 | 0.000 | 0.000 | 0.000 | 0.000 | 38.462 |
| inha | 16.279 | 5.814 | 6.977 | 6.977 | 13.954 | 2.326 | 25.581 | 9.302 | 4.651 | 6.977 | 23.256 | 2.326 |
| pnp | 0.000 | 0.000 | 0.000 | 0.000 | 0.000 | 22.000 | 0.000 | 0.000 | 0.000 | 0.000 | 0.000 | 20.000 |
| sahh | 24.242 | 9.091 | 0.000 | 6.061 | 0.000 | 0.000 | 18.182 | 6.061 | 0.000 | 6.061 | 0.000 | 0.000 |
| tk | 0.000 | 0.000 | 4.545 | 0.000 | 0.000 | 22.727 | 0.000 | 0.000 | 0.000 | 0.000 | 0.000 | 18.182 |
| mean | 7.994 | 9.020 | 4.508 | 10.210 | 11.074 | 14.057 | 9.202 | 10.185 | 5.378 | 12.279 | 14.013 | 16.540 |
| median | 3.964 | 6.143 | 2.532 | 8.432 | 10.296 | 12.947 | 5.267 | 7.353 | 0.000 | 9.801 | 11.732 | 15.800 |

### LIT-PCBA benchmark

Table S18: Proteins chosen from LIT-PCBA as each target representatives.

| | |
|---|---|
| adrb2 | 4lde, 4ldl, 6mxt |
| aldh1 | 4wp7, 5l2o |
| esr1_ago | 1l2i, 2p15, 2qr9 |
| esr1_ant | 1xp1, 2iog, 6chw, 5ufx |
| fen1 | 5fv7 |
| gba | 2v3d, 3rik |
| idh1 | 4i3k, 4umx, 5de1, 5lge, 5sun, 5svf |
| kat2a | 5h84, 5mlj |
| mapk1 | 5v62, 2ojg, 3w55, 4qp3 |
| mtorc1 | 4dri, 4jsx |
| oprk1 | 6b73 |
| pkm2 | 3gqy, 4g1n |
| pparg | 1zgy, 3hod, 3b1m |
| tp53 | 2vuk |
| vdr | 3a2i |

Table S19: ROC AUC, BEDROC, and 5%, 1%, 0.5% EF computed for AutoDock Vina and Convex-PL$^R$ in the LIT-PCBA benchmark

| | ROC AUC | | EF5% | | EF1% | | EF0.5% | | BEDROC$_{\alpha=20}$ | |
|---|---|---|---|---|---|---|---|---|---|---|
| Target | AutoDock Vina | Convex-PL$^R$ | AutoDock Vina | Convex-PL$^R$ | AutoDock Vina | Convex-PL$^R$ | AutoDock Vina | Convex-PL$^R$ | AutoDock Vina | Convex-PL$^R$ |
| adrb2 | 0.373 | 0.506 | 1.176 | 1.176 | 0.000 | 5.880 | 0.000 | 0.000 | 0.042 | 0.090 |
| aldh1 | 0.579 | 0.575 | 1.433 | 1.484 | 1.330 | 1.830 | 1.370 | 2.120 | 0.111 | 0.117 |
| esr1_ago | 0.594 | 0.587 | 1.538 | 1.538 | 7.690 | 0.000 | 0.000 | 0.000 | 0.113 | 0.065 |
| esr1_ant | 0.698 | 0.708 | 2.273 | 2.727 | 3.410 | 3.410 | 6.820 | 2.270 | 0.156 | 0.160 |
| fen1 | 0.519 | 0.499 | 1.167 | 1.944 | 1.110 | 3.890 | 0.560 | 5.000 | 0.053 | 0.088 |
| gba | 0.630 | 0.769 | 2.561 | 5.610 | 5.490 | 17.07 | 6.100 | 25.61 | 0.111 | 0.263 |
| idh1 | 0.614 | 0.551 | 1.026 | 2.051 | 0.000 | 0.000 | 0.000 | 0.000 | 0.049 | 0.089 |
| kat2a | 0.441 | 0.399 | 1.237 | 1.031 | 1.550 | 0.520 | 0.000 | 0.000 | 0.060 | 0.041 |
| mapk1 | 0.647 | 0.642 | 2.208 | 2.013 | 1.950 | 1.950 | 1.950 | 3.250 | 0.108 | 0.095 |
| mtorc1 | 0.438 | 0.452 | 0.619 | 0.825 | 1.030 | 0.000 | 0.000 | 0.000 | 0.033 | 0.043 |
| oprk1 | 0.507 | 0.677 | 0.833 | 0.833 | 0.000 | 0.000 | 0.000 | 0.000 | 0.027 | 0.038 |
| pkm2 | 0.613 | 0.657 | 1.429 | 2.125 | 1.650 | 3.300 | 1.470 | 3.660 | 0.081 | 0.092 |
| pparg | 0.730 | 0.780 | 2.500 | 5.00 | 4.170 | 12.00 | 0.00 | 16.66 | 0.159 | 0.253 |
| tp53 | 0.635 | 0.591 | 3.125 | 1.875 | 3.130 | 4.690 | 3.130 | 3.130 | 0.166 | 0.121 |
| vdr | 0.413 | 0.437 | 0.459 | 0.620 | 0.920 | 0.920 | 0.690 | 0.920 | 0.029 | 0.032 |
| mean | 0.562 | 0.589 | 1.572 | 2.057 | 2.229 | 3.697 | 1.473 | 4.175 | 0.087 | 0.106 |
| median | 0.594 | 0.587 | 1.429 | 1.875 | 1.550 | 1.950 | 0.560 | 2.120 | 0.081 | 0.090 |

### References

(1) Klenin, K. V., Tristram, F., Strunk, T., and Wenzel, W. (2011) Derivatives of molecular surface area and volume: Simple and exact analytical formulas. *J. Comput. Chem. 32*, 2647–2653.

(2) Klenin, K., Tristram, F., Strunk, T., and Wenzel, W. (2012) Achieving Numerical Stability in Analytical Computation of the Molecular Surface and Volume. *From Computational Biophysics to Systems Biology (CBSB11)–Celebrating Harold Scheraga's 90th Birthday 8*, 75.

(3) Karasikov, M., Pagès, G., and Grudinin, S. (2019) Smooth orientation-dependent scoring function for coarse-grained protein quality assessment. *Bioinformatics 35*, 2801–2808.

(4) Kadukova, M., and Grudinin, S. (2017) Convex-PL: a novel knowledge-based potential for protein-ligand interactions deduced from structural databases using convex optimization. *J. Comput.-Aided Mol. Des. 31*, 943–958.
